# Engagement of pulvino-cortical feedforward and feedback pathways in cognitive computations

**DOI:** 10.1101/322560

**Authors:** Jorge Jaramillo, Jorge F. Mejias, Xiao-Jing Wang

**Affiliations:** Center for Neural Science, New York University, New York, NY 10003, USA; Center for Neuroscience, Swammerdam Institute for Life Sciences, University of Amsterdam

**Keywords:** pulvinar, cortex, thalamic reticular nucleus, confidence, hierarchy, working memory, attention, decision making, oscillations, attractor dynamics

## Abstract

Computational modeling of brain mechanisms of cognition has been largely focused on the cortex, but recent experiments have shown that higher-order nuclei of the thalamus, in particular the pulvinar, participate in major cognitive functions and are implicated in psychiatric disorders. Here we show that a pulvino-cortical circuit model, composed of two cortical areas and the pulvinar, captures a range of physiological and behavioral observations related to the macaque pulvinar. Effective connections between the two cortical areas are gated by the pulvinar, allowing the pulvinar to shift the operation regime of these areas during attentional processing and working memory, as well as to resolve decision-making conflict. Furthermore, cortico-pulvinar projections that engage the thalamic reticular nucleus enable the pulvinar to estimate decision-making confidence. Finally, feedforward and feedback pulvino-cortical pathways participate in frequency-dependent inter-areal interactions that modify the relative hierarchical positions of cortical areas. Overall, our model suggests that the pulvinar provides crucial contextual modulation to cortical computations associated with cognition.

## Introduction

The thalamus is involved in a myriad of functions essential to an animal’s survival, including linking the sensory world to the cortex and regulating sleep, alertness, and wakefulness (Ward, 2013). The various thalamic nuclei identified to date form reciprocal connections with the cortex and other subcortical structures (Jones, 2007). Given that abnormal functional connectivity between the thalamus and cortex is a biomarker for psychiatric disorders such as schizophrenia and autism (Anticevic et al., 2014; Nair et al., 2013), it is clinically relevant to understand the neural computations underlying these complex thalamo-cortical loops.

Investigation into the circuit mechanisms of sensory thalamus (Briggs and Usrey, 2009; Petersen, 2007) has already been successful in describing the transformation and the processing of sensory information from the periphery into the cortex. Much less is known about the computations taking place in higher-order thalamic nuclei, i.e., those receiving their driving input from the cortex (Sherman and Guillery, 2013). Far from being a passive relay, the thalamus is now known to play an active role in many of the cognitive functions typically attributed to the cortex alone (McAlonan et al., 2008; Saalmann and Kastner, 2011; Wimmer et al., 2015; Chakraborty et al., 2016; Schmitt et al., 2017).

The primate pulvinar is part of the visual thalamus and is a prominent example of a higher-order nucleus whose exact function remains unresolved (Saalmann and Kastner, 2011; Halassa and Kastner, 2017). Early studies recognized the pulvinar to play a role in attentional processing as single neurons in the pulvinar were modulated by attentional cues (Petersen et al., 1985,1987) and lesions to the pulvinar resulted in attentional deficits including hemispatial neglect towards the contralesional visual field (Wilke et al., 2010, 2013) as well as a deficit in filtering of distractors (Desimone et al., 1990). These results have been confirmed in behavioral and fMRI studies (Danziger et al., 2002), although some of the more subtle effects remain disputed (Strumpf et al., 2013). A recent study showed that the firing rate of neurons in the macaque pulvinar correlated with confidence during a decision-making task with an opt-out component (Komura et al., 2013). Furthermore, calcium imaging of V1-projecting axons of the lateral posterior nucleus (LP), the rodent homologue of the primate pulvinar, revealed that LP signals a mismatch between self-generated and external visual motion (Roth et al., 2015). It is not known how and why the pulvinar contributes to these seemingly disparate cognitive functions.

As part of the visual thalamus, the pulvinar is connected to virtually all of the visual sectors along the cortical hierarchy (Shipp, 2015; Grieve et al., 2000). While the lateral and ventral parts of the pulvinar are connected to early visual cortices (Kaas and Lyon, 2007), the medial pulvinar is connected to association cortices such as the parietal, temporal, and prefrontal cortex (Romanski et al., 1997; Gutierrez et al., 2000). Notably, visual areas and fronto-parietal areas are consistently recruited during tasks that engage or require attention (Buschman and Miller, 2007), working memory (Suzuki and Gottlieb, 2013), and decision-making (Siegel et al., 2015; Hanks et al., 2015). The fact that the neural computations underlying these cognitive functions depend on local, i.e, within-area, as well as on long-range, i.e. across-area, connectivity (Buschman and Kastner, 2015; Christophel et al., 2017; Brody and Hanks, 2016) suggests that the pulvinar could impact cognitive function by modulating cortical computations through pulvino-cortical projections, but a plausible circuit mechanism has not been proposed.

From a physiological point of view, we note that two cortical areas are connected not only via direct, i.e., cortico-cortical, feedforward and feedback projections, but also indirectly via the thalamus (Theyel et al., 2010). It has been hypothesized that these thalamic-mediated indirect projections arising from cortical layer V contribute to the communication between cortical areas (Sherman and Guillery, 2013; Sherman, 2016; Saalmann et al., 2012; Zhou et al., 2016) but the relationship to cognitive function is not clear.

Moreover, cortico-thalamic projections arising from cortical layer VI often engage the thalamic reticular nucleus (TRN), a shell of inhibitory neurons that is an important source of inhibition to the thalamus (Jones, 2007). The TRN-LGN circuit, for example, has been implicated in some forms of attentional control (Wimmer et al., 2015; McAlonan et al., 2008). However, whether the TRN participates in computations related to cognition in tandem with other thalamic nuclei such as the pulvinar remains an open question.

To elucidate the pulvinar’s contributions to cognition, we put forward a framework that connects cortical to thalamic computation. This framework relies on first, a canonical cognitive-type circuit in the cortex (Wang, 2013; Murray et al., 2017), and second, on the existence of two well-defined anatomical pathways that connect the pulvinar to the cortex and back: i) a feedforward or transthalamic pathway that relays cortical information to a second cortical area (Sherman and Guillery, 2013) and ii) a feedback or reciprocal pathway that originates in a given cortical area, targets the TRN and pulvinar, and then projects back to the same cortical area. We built a pulvino-cortical circuit model to map the aforementioned pathways to behaviorally-relevant computations for attention, working memory, and decision making, and to sharpen the interpretation of recent studies that combined pulvinar electrophysiology with behavior (Komura et al., 2013; Saalmann et al., 2012; Zhou et al., 2016). Overall, our results suggest that the pulvinar, through the feedforward and feedback pulvino-cortical pathways, is uniquely positioned to provide crucial contextual modulation to cortical computations associated with cognition.

## Results

### A pulvino-cortical architecture to model two alternative forced choice tasks

We have designed a pulvino-cortical circuit to model cognitive tasks that involve the selection of one of two choices, i.e, two-alternative-forced choice (2AFC) tasks. Such tasks are useful to study distinct components of cognitive processes including attention, working memory, and decision making. The three-module circuit we propose consists of two reciprocally-connected cortical areas and the pulvinar. To model 2AFC tasks, each module consists of two populations of excitatory neurons where each population is selective to one of two stimuli, which can be spatial, directional, or object (Fig. 1). Local connectivity within each cortical module is specified by recurrent excitation and cross-inhibition between the excitatory populations. The local connectivity for each cortical module follows a hierarchical gradient in that the local excitatory recurrence in Module 2 is greater than in Module 1. Long-range connectivity between the two cortical modules is specified by feedforward and feedback projections that are excitatory between same-selectivity populations and inhibitory between different-selectivity populations. Recurrent excitation and cross-inhibition in the two-module cortical circuit can generate winner-take-all dynamics, ramping activity through slow reverberation, and bistability. Thus, the two-module cortical circuit in isolation (i.e., without engagement of the pulvinar) can in principle support a set of cognitive-type computations (Wang, 2013) including visual selection, evidence accumulation during decision-making, and persistent activity for working memory (Murray et al., 2017).

**Figure 1:**
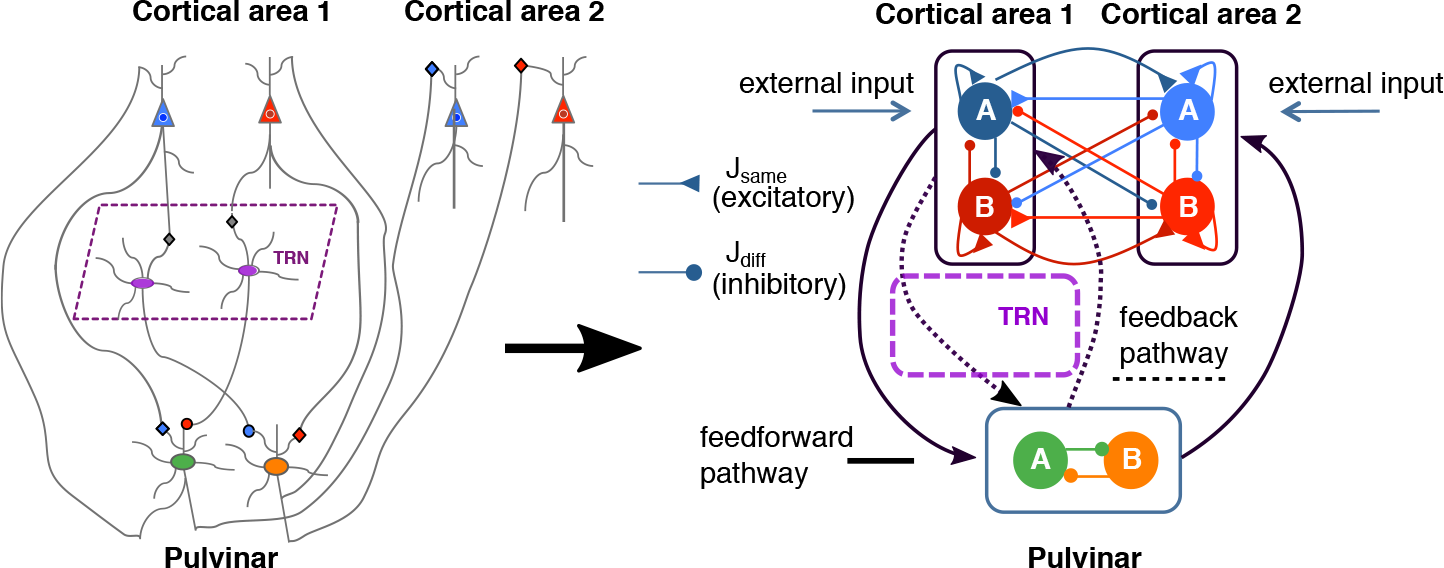
A pulvino-cortical circuit for two-alternative forced choice tasks. The simplified circuit to the right is composed of three modules: two reciprocally-connected cortical modules (1 and 2) and the pulvinar that receives projections from and projects to the cortex through feedforward (solid lines) and feedback (dotted lines) thalamo-cortical pathways. A module here is defined as a set of two excitatory populations (blue and red in cortex, green and orange in pulvinar) where each population is selective to one of two choices, *A* or *B*. In general, synaptic weights *J* can connect two selective populations of either the same (*J*_same_ > 0, excitatory) or opposite (*J*_diff_ < 0, inhibitory) stimulus-selectivity and can be either local (within area) or long-range (across areas). The thalamic reticular nucleus (TRN) allows for long-range disynaptic inhibition from the cortex onto the pulvinar as well as mutual inhibition within the pulvinar. The cortico-pulvino-cortical connections follow the general topography of the cortico-cortical connections. Synapses labeled with triangles and circles denote effective excitatory and inhibitory connections, respectively.

To establish the connectivity between the pulvinar and the two cortical areas in our model, we distinguish two pathways (Jones, 2007; Sherman and Guillery, 2013): i) a transthalamic feedforward pathway that includes a projection from cortical area 1 to a pulvinar relay cell population followed by a projection from the aforementioned relay cells to cortical area 2 and ii) a feedback pathway that originates in either of the cortical areas, targets the thalamic reticular nucleus (TRN) and pulvinar, and then projects back to the same cortical area. The TRN allows for cortical disynaptic inhibition onto pulvinar cells and, because of a lack of excitatory recurrency within the pulvinar (Jones, 2007), mutual inhibition between pulvinar cells. We will examine explicit TRN-pulvinar projections at a later stage when we discuss the cortico-thalamic feedback circuit in detail. Overall, the pulvino-cortical projections in our model follow the topography of the cortico-cortical projections: excitatory between same-selectivity populations and inhibitory between different-selectivity populations.

For the tasks modeled in Figs 2-6, we will study the type of information represented in the pulvinar, and how this information modulates the cognitive-type computations in the cortex via the feedforward and feedback pulvino-cortical pathways. We will also consider other functional topologies in Fig. S1 and a circuit with laminar structure in Fig. 7 when we discuss frequency-dependent inter-areal interactions.

**Figure 2:**
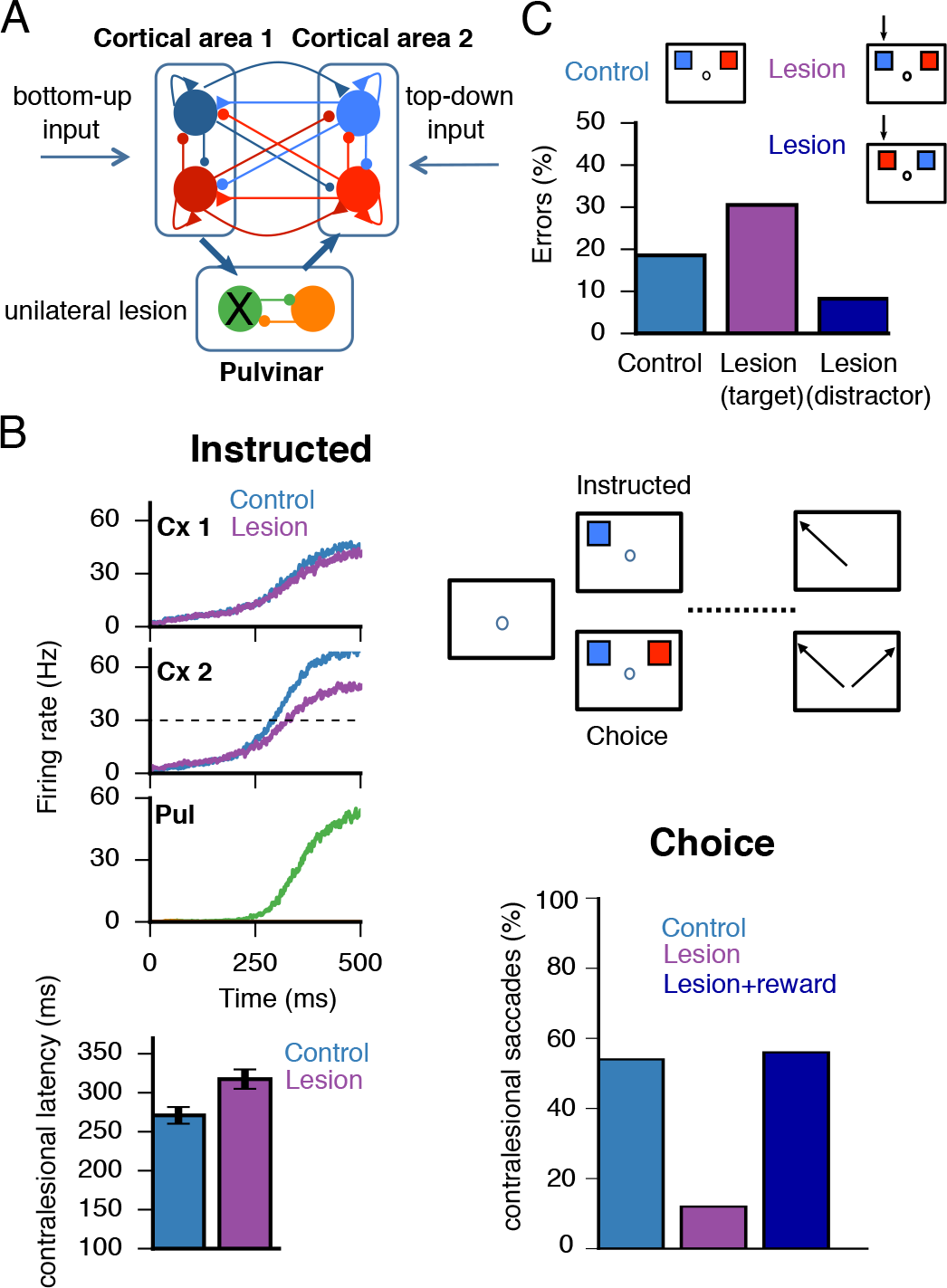
Pulvinar lesion-induced gain imbalance produces asymmetric attentional deficits. ***A,*** Schematic as in Fig. 1 where external inputs are labeled as either bottom-up (sensory) or top-down (internal), with pulvinar excitability λ =230 Hz/nA. A unilateral lesion is shown that affects the left visual field. Topography thus corresponds to visual and not anatomical space. ***B,*** Visuospatial task based on Wilke et al. (2013), where a subject must make a saccade towards a visual target after a delay period (instructed) or select one of two simultaneously presented visual targets on opposite sides of the visual field (choice). In the instructed task, saccade latencies towards the contralesional field are larger than in controls. In the choice task, the proportion of saccades to the contralesional field is reduced compared to controls, but ameliorated with the addition of reward (Wilke et al., 2013). ***C,*** Visuospatial task modeled after Desimone et al. (1990) where a subject must attend to and select a target (blue) that was flashed at the same position as a cue presented during fixation. A distractor (red) is presented simultaneously in the opposite hemifield. Simulations are performed for control and unilateral lesion of the pulvinar. Black arrows point to the affected visual hemifield and two conditions can be distinguished: either the target (magenta) or the distractor (dark blue) lies within the affected hemifield. Error rates shown below.

**Figure 3:**
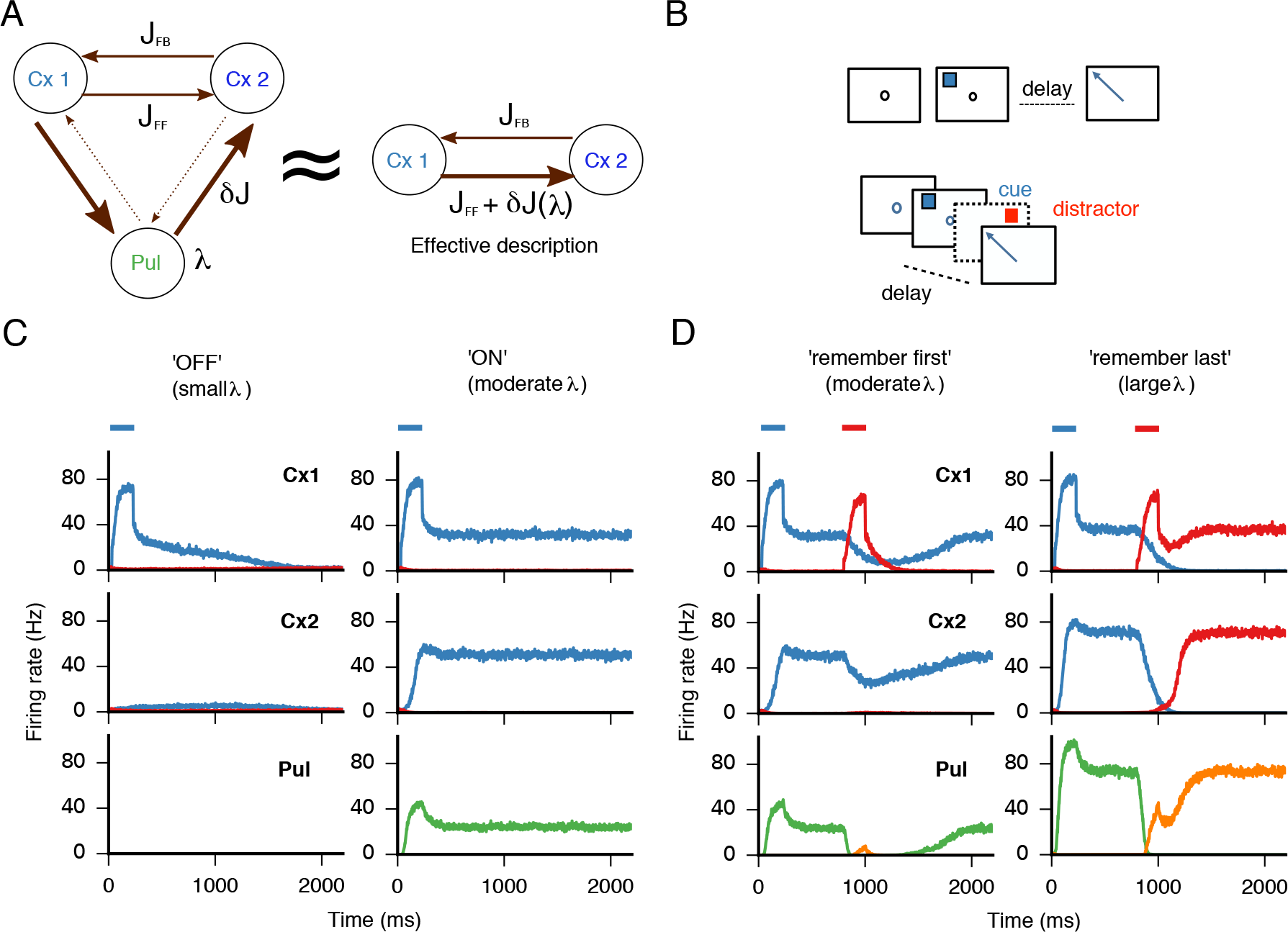
Gating of effective cortico-cortical connectivity and persistent activity through pulvinar gain modulation. ***A***, A three-module pulvino-cortical architecture is equivalent to a two-module cortical architecture, where the effective cortico-cortical connectivity is controllable via the pulvinar excitability parameter λ, and δJ denotes the λ-dependent extra connectivity provided by the transthalamic route. ***B,*** Schematics of the tasks in *C* (top, simple memory-saccade task) and *D* (bottom, memory saccade with distractor during the delay period) ***C,*** In a simple memory saccade task, persistent activity in the cortico-thalamic system is contingent on the activation of the pulvinar, which can act as a switch. When the pulvinar is ‘off’ (λ = 120 Hz/nA), the activity decays in the first cortical area and no activity is observed in the rest of the pulvino-cortical system. When the pulvinar is ‘on’ (λ = 220 Hz/nA), reciprocal loops with the cortex are enough to sustain reverberant activity in the cortico-thalamic circuit, and a global attractor is reached. ***D,*** In a memory-saccade task with a distractor, the pulvinar can control the response of the system by biasing the circuit into making the system more (‘remember first’, λ = 220 Hz/nA) or less (‘remember last’, λ = 290 Hz/nA) robust to distractor interference. Blue and red bars denote target and distractor presentation times, respectively.

**Figure 4:**
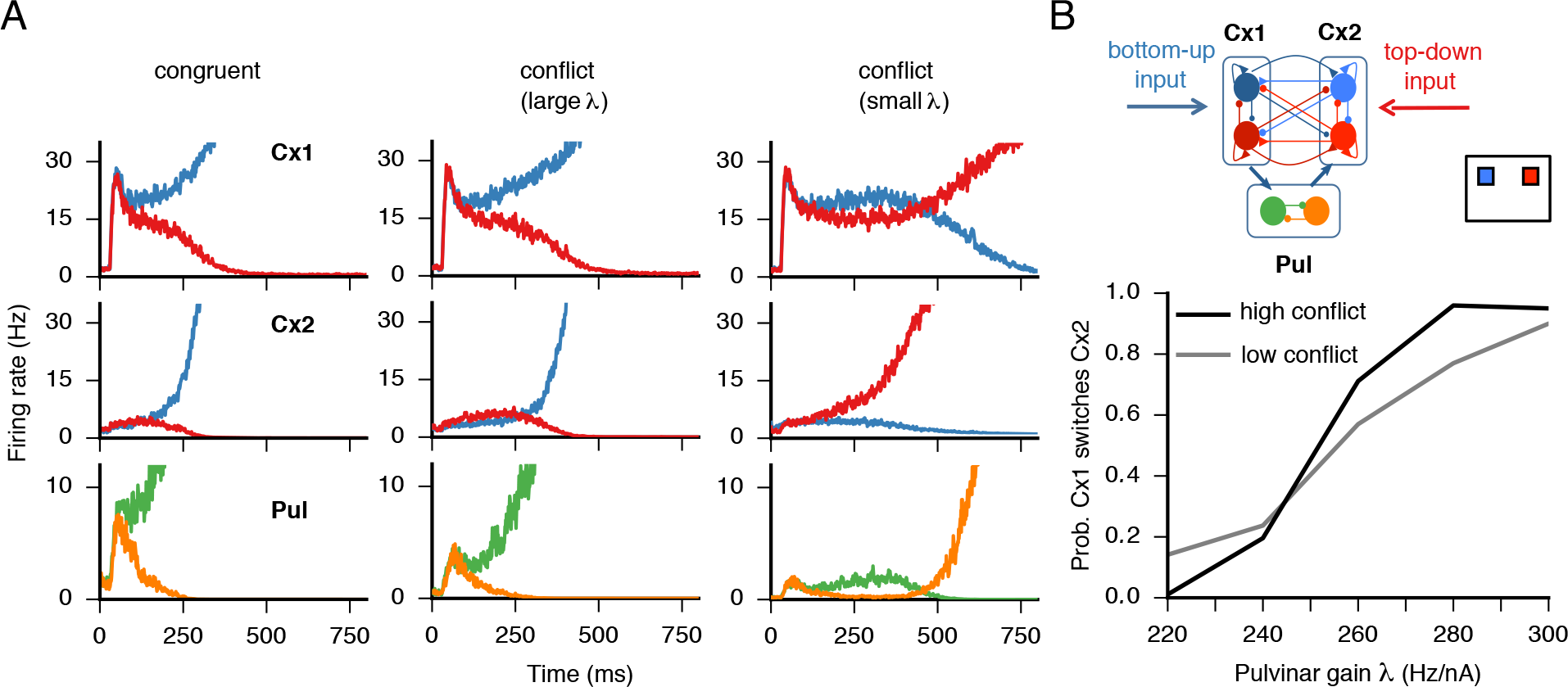
Pulvinar-mediated effective connectivity between cortical areas resolves decision-making conflict. ***A,*** Conflict resolution in the pulvino-cortical model. In the congruent scenario (left), bottom-up and top-down inputs target populations with the same selectivity so that a consistent decision is made. In the conflict scenario (middle and right), bottom-up input favors the blue excitatory population in cortical area 1 while top-down favors the red excitatory population in cortical area 2, resulting in inter-areal competition. For large λ (λ = 280 Hz/nA), the effective feedforward pathway connecting cortical area 1 to 2 is preferentially biased so that the choice reflects bottom-up information (middle). For small λ (λ = 220 Hz/nA), the effective feedforward strength is decreased so that the choice reflects top-down input (right). High (*c*′ = 20) and low (*c*′ = 10) conflict trials are shown in thick and thin lines, respectively. ***B,*** Schematic of conflicting stimuli and responses in the pulvino-cortical circuit (top). In the conflict scenario, the probability of cortical area 1- bottom-up recipient-enforcing its encoding to cortical area 2 - top-down recipient-depends on the value of the pulvinar excitability λ and on the conflict level *c*′ (bottom).

**Figure 5:**
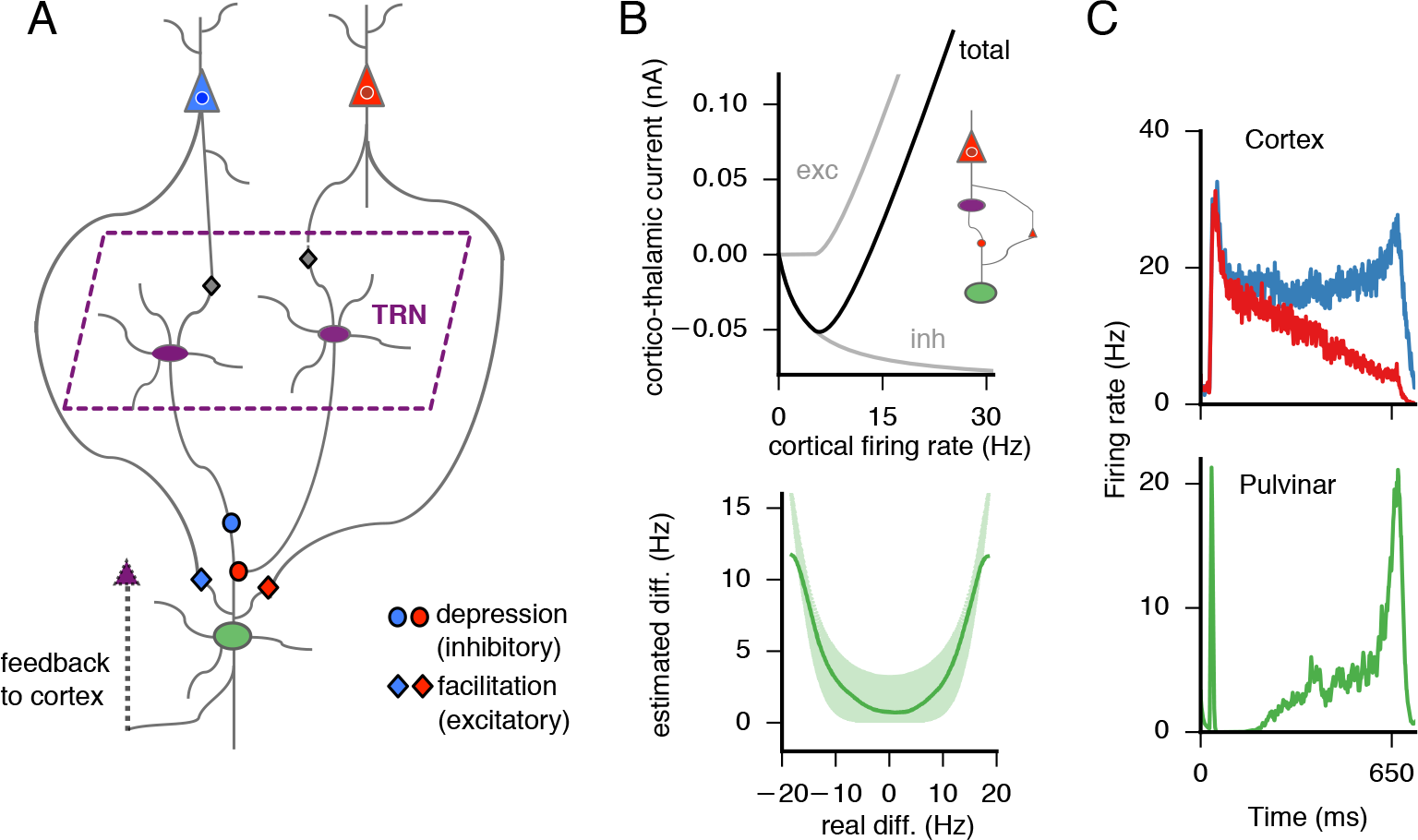
Calculation of absolute differences by a circuit that engages the cortex, pulvinar, and TRN. ***A,*** The cortical circuit component consists of two cortical populations (here schematically represented by single neurons) differentially selective to two distinct stimuli, blue and red. As in Fig. 1, the cortical excitatory populations receive lateral projections and interact through a common pool of interneurons (connections are not shown for clarity). The two cortical populations are connected to a pulvinar cell via a direct excitatory monosynaptic connection and a disynaptic inhibitory projection. The excitatory connection exhibits short-term facilitation while the inhibitory TRN-pulvinar connection exhibits short-term depression. ***B,*** Top, the short-term synaptic dynamics in the thalamo-cortical circuit result in non-linear function of the cortical firing rate so that the input is effectively inhibitory for low firing rates but excitatory for high firing rates. Inset shows the motif that generates the plot for a single cortical cell. Bottom, if the activity of both cortical cells is combined, the resulting activity at the level of the pulvinar (λ =300 Hz/nA) resembles approximately an absolute value function of the difference between the firing-rate activities of the two cortical cells. ***C,*** Firing activites of the cortex (top) and pulvinar (bottom), where the pulvinar integrates the cortical activity and approximately calculates the absolute value of the difference between the activities of the competing cortical populations.

**Figure 6:**
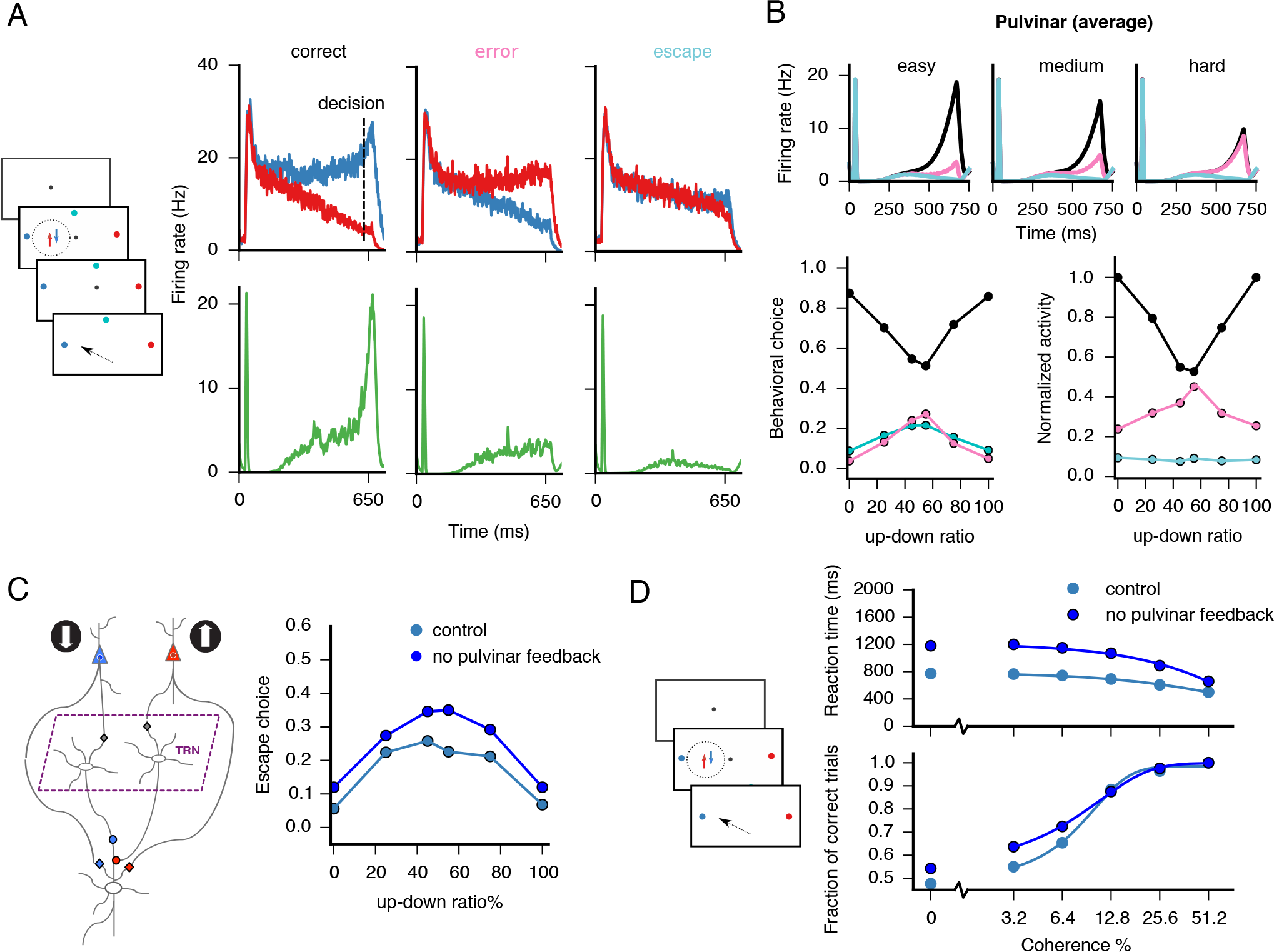
A pulvino-cortical circuit for estimating decision-making confidence. ***A,*** schematic oi the task is shown on the left (see details in main text). Single cells in the pulvinar (green, bottom) represent confidence through their firing rate for correct, error, and escape trials. A necessary condition for a correct trial is that the cortical population representing more evidence, here the blue population (top), has a greater activity than the population representing less evidence, the red population, at the time of decision. Moreover, for both correct and error trials, the difference between the activities at the decision time must be greater than a predefined bound ϵ = 4 Hz. Otherwise, the subject forgoes the decision and escapes, (opts out). ***B,*** Top, average pulvinar firing rates as a function of difficulty (easy, medium, hard) and trial type (correct,black; error,pink; escape,cyan), color coded as in *A*. Bottom, behavioral choice (left) and normalized pulvinar activities (right) as a function of difficulty and trial type. ***C,*** Simulated unilateral lesion to the pulvinar, i.e., no feedback to the cortex, causes an increase in escape frequency with respect to control. ***D,*** In a reaction-time version of the random-dot discrimination task, a lesion to the pulvinar causes a speed-accuracy tradeoff, more noticeable at low coherence levels.

**Figure 7:**
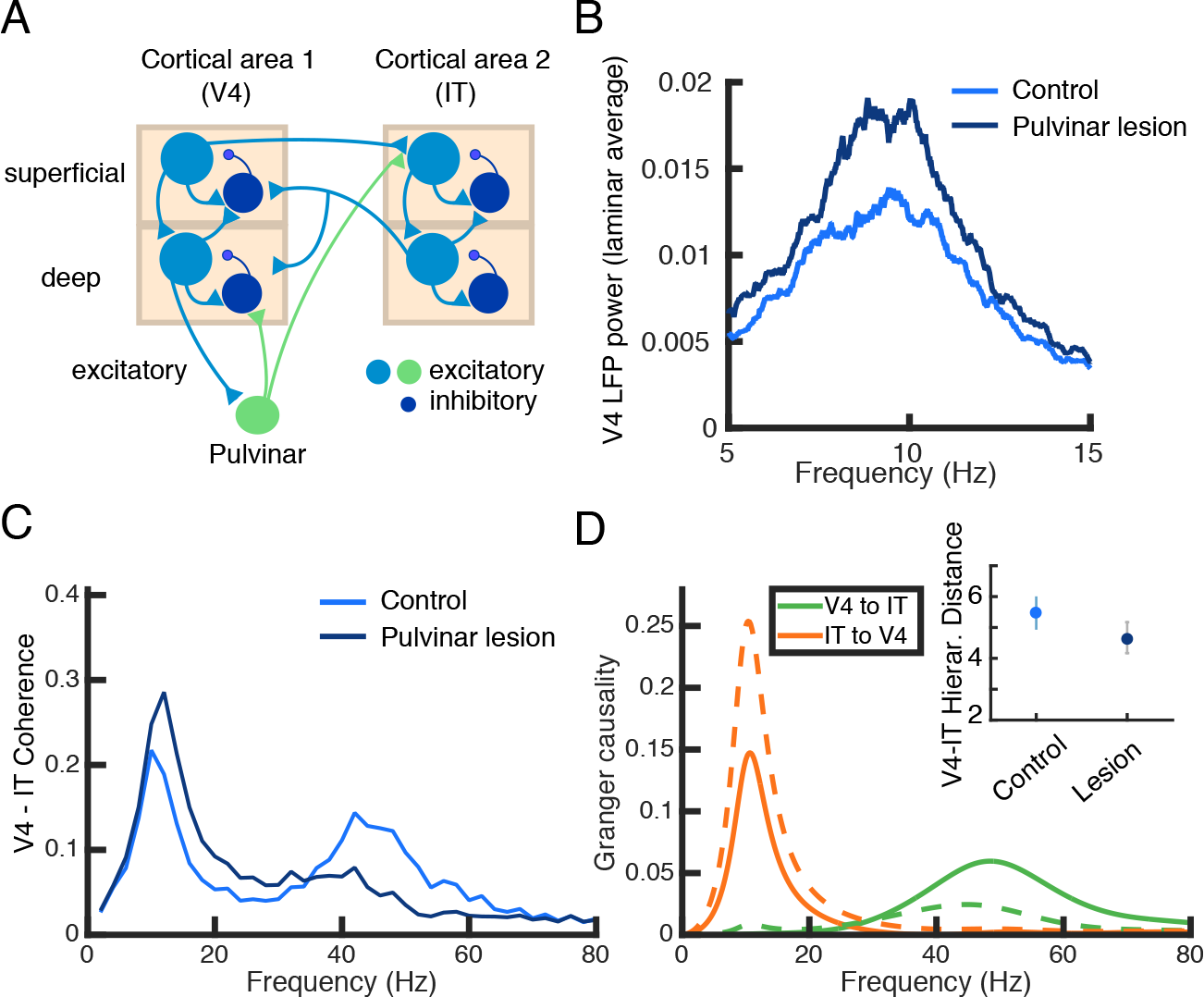
Thalamic gating of gamma and alpha oscillations across cortical areas (see also Zhou et al. (2016); Saalmann et al. (2012)). ***A,*** Schematic of a distributed pulvino-cortical circuit with laminar structure. The model is composed of two reciprocally connected cortical modules (here V4 and IT) and the pulvinar that both receives projections and projects to the cortical modules. The transthalamic projection targets layer IV in the cortical area 2, which is then relayed to the superficial layers. ***B,*** After a lesion to the pulvinar, the power measured from the V4 population activity exhibits an increase in low-frequency oscillations. ***C,*** The two cortical areas are coherent at gamma frequencies and this coherence is decreased after lesioning the pulvinar. ***D,*** The coherence effects observed in C extend to Granger causality, which in addition measures directionality. Control and pulvinar lesion scenarios are shown in solid and dashed lines, respectively. Inset shows that the hierarchical distance between the cortical areas decreases after a pulvinar lesion.

### Pulvinar lesion-induced gain imbalance produces asymmetric attentional deficits

To better understand and constrain the dynamics of our pulvino-cortical circuit, we first examine the impact of unilateral lesions to the pulvinar. Indeed, lesion studies have provided important insights into the role of the pulvinar in tasks that engage attention (Wilke et al., 2010, 2013; Snow et al., 2009; Desimone et al., 1990). At least two distinct effects have been observed after unilateral lesions of the pulvinar: hemispatial neglect, whereby one area of the visual field is unaccessible either due to lack of perceptual awareness or motivation (Wilke et al., 2010, 2013) and a deficit in distractor filtering, whereby performance in a visual search task decreases when a target is flanked by irrelevant distractors (Desimone et al., 1990; Fischer and Whitney, 2012; Snow et al., 2009; Strumpf et al., 2013). We used the pulvino-cortical architecture introduced above to model the behavioral effects after a unilateral lesion of the pulvinar and, more generally, to elucidate the computations in the pulvinar (Fig. 2).

The first task was modeled after Wilke et al. (2013). In this task, subjects have to select a target that appears on a screen after a fixation period. In the Instructed variant of the task, only one target is presented and subjects have to make a saccade towards the cued target to obtain a reward. In the Choice variant, the subjects are presented with two targets that yield equal reward when selected. We modeled the contrast of the targets, a bottom-up input, as an input current to the first module and modeled reward expectation, a top-down input, as an input current to the second cortical area (see Figs. 2 *A, B* and Methods).

We modeled a lesion by fixing the firing rate of one of the pulvinar populations to zero. The basic finding is that after the lesion there is an attentional disruption in the contralesional field of lesioned subjects with respect to control (Fig. 2 *B*). On Instructed trials, unilateral lesions cause an increase in saccade latency towards the contralesional field (see also Tanaka (2006)). On Choice trials, the proportion of saccades towards the contralesional field decreases as compared to control. Interestingly, this effect is ameliorated by the addition of more reward to the target on the contralesional side, as reported by Wilke et al. (2013). In our model, such attentional deficits are observed because the lesion effectively reduces the excitation towards the contralesional, i.e., affected visual hemifield, which in turn induces a gain imbalance in the multi-regional circuit. This pulvinar-induced imbalance is quickly amplified by the recurrent circuitry in the cortex and propagated asymmetrically throughout the pulvino-cortical circuit to produce the impairments described.

Pulvinar lesions are known to affect distractor processing in humans and non-human primates (Desimone et al., 1990; Danziger et al., 2002; Snow et al., 2009). To understand why this is the case, we modeled a second task after Desimone et al. (1990) where a subject must attend to and select a target that was flashed at the same position as a cue presented during fixation (Fig. 2*C*). A distractor was defined as another stimulus simultaneously flashed at an opposite location to the target (Desimone et al., 1990). In our model, the behavioral relevance of the target is associated with a larger value of the differential input *c′* (Eq. 13) which represents the target-distractor similarity, i.e., low values of *c*′ represent difficult trials. We found that only when the target was located in the affected visual hemifield (opposite to the site of the simulated anatomical lesion), the error rate increased with respect to controls. A slight improvement in performance was observed in the opposite scenario, when the distractor was located in the affected hemifield (Wilke et al., 2010; Desimone et al., 1990). In essence, the non-linear winner-take-all circuit effectively suppresses representations that are not as behaviorally relevant as the target. Along these lines we suggest that the topography of the pulvino-cortical connections, i.e., excitatory projections between cells having similar selectivity and cross-inhibition between cells with opposite selectivity, is the structural mechanism underlying distractor filtering.

It’s important to note that we used the lesion vs control simulations to set the basic parameters for the cortical and thalamic modules that will be used in the rest of the figures (see Table 1).

**Table 1.**
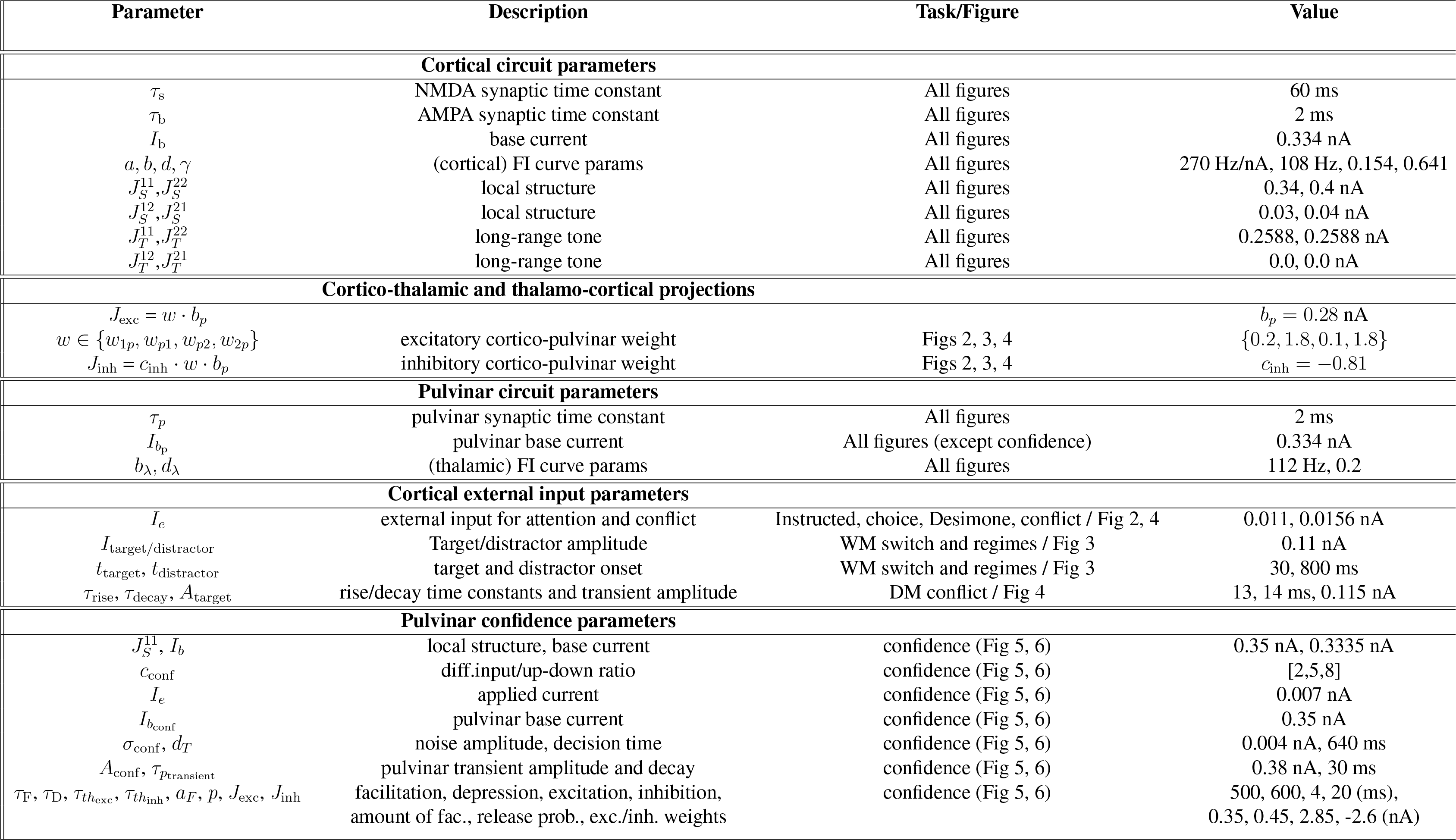
Parameters for numerical simulations

### Gain modulation in the pulvinar flexibly controls effective cortico-cortical connectivity

The model of simulated lesions described above hints at a generalized gain function for the pulvinar (Purushothaman et al., 2012) that can potentially impact cortical processing and behavior. Given that our distributed circuit model includes direct cortico-cortical projections as well as indirect transthalamic projections, we examined what the hypothesized gain function of the pulvinar implies for cortical processing. In our model, two cortical areas are reciprocally connected via direct anatomical projections, but also indirectly connected through interactions with the pulvinar. Therefore, the total connectivity between the two cortical areas - here referred to as “effective” connectivity - has two contributions: a direct cortico-cortical projection and an indirect projection provided by the transthalamic route that engages the pulvinar. We can show that the amount of extra connectivity from the transthalamic route depends on the pulvinar excitability λ, here defined as the slope of the input output FI curve in the pulvinar (see Methods, Eq. 4). Indeed, the three-module pulvino-cortical system is approximately equivalent to a two-module cortico-cortical system with a new effective connectivity matrix (Fig. 3*A* and Eq.14). This new effective connectivity matrix depends on important parameters such as the pulvinar time constant *τ*_*p*_, and more importantly, the pulvinar excitability λ. In particular, if we assume that the feedforward relay weights in the hierarchy-preserving direction (cortical area 1 → pulvinar → cortical area 2) is larger than in the reverse direction,the overall feedforward strength between the two cortical areas can be modulated by the pulvinar excitability λ, with the feedforward strength growing linearly with λ (see Eq. 20). The fact that the effective connectivity between two cortical areas depends on the pulvinar excitability λ, means that such effective connectivity is controllable via external input onto the pulvinar, for example, via top-down modulation from prefrontal cortex (Zikopoulos and Barbas, 2006; Romanski et al., 1997) or superior colliculus (Baldwin et al., 2013). Notably, this form of open-loop control does not depend on any oscillatory mechanisms (Saalmann and Kastner, 2009), although we will later show that gating of cortical oscillations (Zhou et al., 2016; Saalmann et al., 2012) is readily achievable.

In the following we examine the computational implications of the pulvinar-mediated effective connectivity between two cortical areas in the context of working memory and decision-making tasks.

### Pulvinar-mediated gating of persistent activity in the pulvino-cortical circuit

Spatially-selective persistent activity is another cognitive computation that is subserved by the cortex, possibly across multiple cortical areas (Suzuki and Gottlieb, 2013; Christophel et al., 2017). Here we examine how the pulvino-cortical circuit can sustain spatially-selective persistent activity in a distributed fashion (Fig. 3*B* - *D*). We assume that the pulvinar is subject to top-down control such that its excitability (here represented by λ) is variable and potentially a function of behavioral state. We examine how the pulvinar-induced modulated connectivity between two cortical areas affects working-memory computations in the pulvino-cortical circuit.

In Figure 3*C*, we model a simple memory task where a stimulus is presented briefly, and the subject must remember the location of the stimulus as typically done in attentional cuing (Saalmann et al., 2012) and/or memory-saccade tasks (Wilke et al., 2013; Suzuki and Gottlieb, 2013). We consider two scenarios corresponding to two values of the pulvinar excitability λ: a “small” and “moderate” value of λ. If λ is small (pulvinar ‘off’), the pulvinar is not actively engaged and the distributed circuit cannot reach a global persistent state: the activity of cortical area 1 decays after vigorously responding to the transient stimulus. In this case, there is no propagation to the second cortical area (Theyel et al., 2010) and the excitatory recurrency in cortical area 1 is not sufficient to sustain a persistent-activity (attractor) state. On the other hand, for a larger value of λ (pulvinar ‘on’) the circuit can reach a state in which both cortical areas and the pulvinar exhibit spatially-selective persistent activity. In this case, the pulvinar effectively augments long-range projections that help sustain a persistent-activity state in the pulvino-cortical circuit - a global attractor-even if the cortical circuits do not exhibit persistent activity independently (Murray et al., 2017). To conclude, our results show that the pulvinar can act as a λ-controlled memory switch. These results are consistent with the engagement of the pulvinar in visuo-spatial tasks (LaBerge and Buchsbaum, 1990), in particular when a transient cue induces persistent activity in the pulvinar (Saalmann et al., 2012; Halassa and Kastner, 2017). Along these lines, we suggest that the documented involvement of various thalamic nuclei in delay tasks (Mitchell and Chakraborty, 2013; Schmitt et al., 2017; Guo et al., 2017; Bolkan et al., 2017) extends to the pulvinar.

We also analyze the behavior of the distributed pulvino-cortical circuit with respect to temporal processing in a memory-saccade task with distractors (Figure 3*D*). A distractor is operationally defined as a stimulus presented during the delay period after the target, but otherwise identical in amplitude and duration (Suzuki and Gottlieb, 2013). Again we consider two values of the pulvinar excitability λ, “moderate” and “large”. Similar to the scenario considered in Figure 3*C*, the circuit is able to sustain a spatially-selective memory state given a sufficiently large value of λ. The behavior of the circuit with respect to distractor processing, however, will depend on how large λ is. If the value of λ is moderate (Figure 3*D*, left), there is propagation to the second cortical area and the extra feed-forward synaptic connectivity is moderately engaged. In this regime, there is enough feedforward drive to engage cortical area 2 to help sustain a more stable attractor and the response to the distractor becomes smaller and transient, especially in cortical area 2 (Murray et al., 2017). On the other hand, if the value of λ is large enough (Figure 3*D*, right), the extra feedforward synaptic connectivity will be markedly engaged, causing the incoming distractor input to be more effectively propagated to cortical area 2. Thus, the strong engagement of the distractor is enough to override the encoding of the target in this regime.

We suggest that the distributed pulvino-cortical circuit model can operate in two regimes, depending on the value of the pulvinar excitability λ: a ‘remember-first’ regime if λ is moderate, and a ‘remember-last’ regime, when λ is large. The former scenario is consistent with the reported differences in distractor processing between LIP and pre-frontal cortex (cortical areas 1 and 2 in the model, respectively) during a working memory task (Suzuki and Gottlieb, 2013). The latter scenario is consistent with pulvinar involvement during distractor-induced interruption of goal-oriented tasks (Michael et al., 2001) (see also Bisley and Goldberg (2006) for analogous results in LIP). To summarize, our model suggests that the transthalamic feedforward pathway allows the pulvino-cortical cognitive circuit to operate in two distinct working memory regimes, thus augmenting the computational capabilities of an otherwise isolated cortical circuit with fixed long-range connectivity.

### Pulvinar-mediated effective connectivity between cortical areas resolves decisionmaking conflict

Decision making is a cognitive function that potentially involves multiple areas (Komura et al., 2013; Buschman and Kastner, 2015; Brody and Hanks, 2016; Hanks et al., 2015). We explore the relationship between the pulvinar-mediated effective connectivity modulation introduced above and decision making. We examine the functioning of the pulvinar-cortical circuit where, as before, the cortical modules are endowed with a winner-take-all mechanism and can accumulate sensory evidence due to the slow NMDA-receptor dynamics. In particular, we consider a conflict scenario, whereby bottom-up and top-down inputs compete for attention and selection to two stimuli located on opposite sides of the visual field (Figure 4). This scenario could result from, for example, a competition between a bottom-up signal such as luminance biasing one side of the visual hemifield and a top-down signal such as reward expectation biasing the opposite hemifield during visual selection (Markowitz et al., 2011). To model such conflict scenario, we consider external inputs to the circuit that can be segregated into “bottom-up”, targeting cortical area 1 and “top-down”, targeting cortical area 2, following hierarchical processing (Buschman and Miller, 2007). Our pulvino-cortical circuit model predicts that when the pulvinar excitability λ is large, the effective feedforward pathway from cortical area 1 to 2 is strengthened, so that ultimately the choice within cortical area 1 is represented in the pulvino-cortical system (Figure 4 *A*, middle). In contrast, when the pulvinar excitability λ is small, the effective feedforward strength is small (Figure 4 *A*, right) and cortico-cortical feedback enables the choice within cortical area 2 to be represented in the pulvino-cortical system. The conflict scenario modeled in Fig. 4 receives support from a fMRI study from Rotshtein et al. (2011) who showed that pulvinar resolves the competition between working memory (WM) and visual search: the WM process interfered with the visual search as if the recalled WM item were a distractor. Importantly, the WM-induced distraction in Rotshtein et al. (2011) was accompanied by a decrease in pulvinar activity with respect to control, as hypothesized by our model with small λ (Figure 4 *A*, right). To conclude, our results suggest that the pulvinar mediates the competition between modules or processes across cortical areas that complements the competition between features-here spatial locations-within a cortical area.

In Figure 4 *B* we show that the probability of cortical area 1 (bottom-up input recipient) enforcing its choice on cortical area 2 (top-down input recipient) increases as a function of the pulvinar excitability λ. In the case of high conflict between bottom-up and top-down stimuli (high value of *c*′), the transition to switching cortical area 2 is more abrupt as compared to the case of low conflict. Overall, we suggest that gain modulation in the pulvinar can resolve cortical competition and the outcome of such competition depends on the externally-controlled pulvinar gain.

### A cortico-TRN-pulvinar circuit can account for the decision-making confidence signals observed in pulvinar

In the sections above we have examined some of the computational capabilities of the transthalamic route that indirectly connects two cortical areas. Now we analyze the cortico-thalamo-cortical feedback pathway more closely and examine why such pathway might be related to the representation of confidence in the pulvinar in the context of decision making (Komura et al., 2013).

In this study we refer to the confidence concept in the sense of decision-making confidence: the subjective probability or belief that the chosen option is correct based on the evidence contributing to it (Kepecs et al., 2008; Hangya et al., 2016; Pouget et al., 2016). In a landmark study - and particularly relevant to our computational model- Kiani and Shadlen (2009) observed that during a decision-making task (the Kiani task), both the decision and the confidence associated to that decision was related to cortical activity in area LIP of the macaque. In the Kiani task, decision confidence in particular could be assessed due to the task design that included an opt-out component: the subject had the option to either make a decision based on the stream of evidence and obtain a sizable reward if correct or, conversely, opt out to obtain a smaller reward. For correct trials, the accumulation of sensory evidence eventually led to ramping activity of a population of neurons within their choice receptive field, thus reflecting a decision (Kiani and Shadlen, 2009; Roitman and Shadlen, 2002). For trials where the subject opts out, however, the firing rates of neurons both within and outside their receptive field reached intermediate levels. Thus the subject was more confident, i.e., would opt out less often, when there was a relative divergence of LIP activity during choice behavior. More precisely, the difference between the firing rate traces within and outside the response field predicted a confidence level(Wei and Wang, 2015).

In a related study, Komura et al. (2013) found single neurons in the medial pulvinar of the macaque whose firing rate predicted whether the animal, in another version of an opt-out task (Komura task), would opt out. In contrast to the LIP neurons in the Kiani study, pulvinar neurons in the Komura study represented confidence explicitly: a single firing-rate trace was informative of the confidence level. The characterization of decision-making confidence in the Kiani and Komura tasks prompts the following question: why do pulvinar cells represent confidence via their firing rate and how is this representation related to the implicit confidence representation in cortex? Given the known connectivity between parietal cortex and pulvinar (Gutierrez et al., 2000), we explored how a cortico-thalamo-cortical feedback pathway could contribute to the representation of decision confidence in the pulvinar.

We propose a pulvino-cortical circuit model to elucidate the mechanisms behind confidence-related computations in cortex and pulvinar. The reciprocally connected pulvino-cortical circuit is based on that of Fig. 1 but now contains explicit TRN-pulvinar connections as shown in Figure 5 *A*. For simplicity, we focus on one cortical module. The cortical module is composed of two excitatory populations that are selective to two stimuli *A*, *B* (e.g., motion direction). The cortical populations exhibit winner-take-all dynamics and can accumulate sensory evidence. The cortical module sends projections to the pulvinar that first target the TRN and receives thalamo-cortical feedback in return. We first describe this connectivity in more detail and then analyze how this connectivity relates to pulvinar activity during a decision making task.

We assume that a model pulvinar neuron integrates input from two excitatory populations from the same cortical area. Such cortico-thalamic input includes first, a direct monosynaptic connection from cortex to pulvinar and second, an indirect disynaptic connection through the TRN. In our model, there is a pair of such connections for both cortical populations selective to *A* and *B*, respectively (Figure 5 *A*). Crandall et al. (2015) has shown in slice physiology experiments that the direct excitatory cortico-thalamic projection in the somatosensory thalamus exhibits short-term facilitation while the inhibitory TRN-pulvinar projection exhibits short-term depression (see also Kirchgessner and Callaway for similar results in rodent pulvinar, in-vivo, SfN Abstract 2017). We analyze the implications of these plastic projections in the macaque pulvinar in-vivo during a decision-making task.

During the sensory evidence accumulation process, the cortical populations *A* and *B* compete for a choice resulting in a “winner” (for example, *A*) whose firing rate ramps up while the “loser” population (for example, *B*) ramps down (Figure 5 *C*, top). In this scenario, when the firing rate *r*_*A*_ of cortical population *A* is high, the direct cortico-pulvinar excitatory synapse - from cortical population *A* to pulvinar-facilitates while the respective inhibitory TRN-pulvinar synapse depresses. This results in a net positive current from population *A* to the pulvinar (Figure 5 *B*, top). Due to competition between the populations *A* and *B* during decision-making, the firing rate *r*_*B*_ would be low in this scenario, and neither the direct cortico-pulvinar excitatory synapse - from cortical population *B* to pulvinar-facilitates nor the respective inhibitory TRN-pulvinar synapse depresses. Thus the strong TRN-pulvinar connection results in an effective negative current from population B to the pulvinar (Figure 5 *B*, top). Overall, the positive and negative contributions from the cortical activity result in a cortico-pulvinar current that approximately scales as *r*_*A*_ − *r*_*B*_. Since the pulvino-cortical circuit is symmetric, *r*_*B*_ − *r*_*A*_ will also be represented in case population *B* wins the competition. We can therefore show that the pulvino-cortical circuit approximately calculates |*r*_*A*_ − *r*_*B*_|, i.e., the pulvinar represents the absolute value of the difference of the activities between the two afferent cortical populations, (see Figure 5 *B*, bottom, and Methods after Eq. 22 for details of the calculation). Thus, the stimulus-selective cortical activity in the cortex is effectively transformed to non-selective differential activity in the pulvinar via the plastic cortico-thalamic projections that engage the pulvinar and the TRN (Fig. 5*C*).

Now we study the implications of the pulvino-cortical circuit model in the context of a decision-making task with an opt-out component. We first consider a fixed-duration version of the task (Figure 6 *A*; see also Komura et al. (2013); Kiani and Shadlen (2009)) where the subject is presented with a display of random dots and has to decide on the net-direction of motion of the display for varying levels of difficulty. Crucially, the subject has the option to forgo the sensory-based decision and opt-out - referred to as ‘escape’ by Komura et al. (2013) - for a smaller but sure reward. We modeled such a task by considering motion-direction selective inputs to two cortical populations (Figure 5*A* and 6*A*). Due to the trial-to-trial stochastic nature of the cortical response to the stimulus (Eq. 5), the cortico-thalamic circuit model can reproduce correct and error trials, as well as escape trials for which the cortical activities have not diverged (Kiani and Shadlen (2009) and Figure 6 *A*; see figure caption for details of the different trial types). Furthermore, the decision-making readout in the cortex results in a specific psychophysical performance: for correct trials the proportion of choices exhibits a V-shape as a function of task-difficulty (inverse to the coherence in Roitman and Shadlen (2002), see Eq. 29) while the opposite is true for error and escape trials (Figure 6 *B*, bottom left). Concurrent with the cortical-based read-out of the decision, the pulvinar integrates the activity of the two populations and calculates an approximate absolute value of the cortical firing rate differences, as in Fig. 5*B*. We found that pulvinar responses signaled via their firing-rate amplitude whether a given trial was correct, error, or escape (Figure 6 *B*, top and bottom right). We suggest that if a cortical area (e.g., parietal cortex) represents decision-making confidence via the activities of two neural populations (an implicit representation of confidence; Kiani and Shadlen (2009); Wei and Wang (2015)), the plastic pulvinar-TRN circuitry will transform the implicit representation of confidence in the cortex to an explicit representation in the pulvinar (Komura et al. (2013), their figure 3).

We tested the role of the return projection from the pulvinar to the cortex (‘feedback to cortex’ in Fig. 5*A*) by simulating a lesion to the pulvinar. We found that after the lesion, the number of escape responses increased with respect to control, notably for low-coherence, i.e., difficult, trials (Figure 6 *C*), also consistent with the Komura et al. (2013) study. Indeed, we assumed that the return projection targeted both selective populations in the cortex equally and a pulvinar lesion reduces the input drive of the winner-take-all mechanism in the cortical module (Wong and Wang, 2006). We also simulated a reaction-time version of the task without an opt-out component in control and pulvinar-lesion scenarios. We found a speed-accuracy tradeoff: the circuit with the lesioned pulvinar exhibited slower but slightly more accurate responses (Figure 6 *D*). Indeed, a lesion in the pulvinar reduces the overall excitation in the cortex which makes the decision-process slower by giving the system more time to integrate information, which in turn slightly improves performance for difficult trials. We contend that the pulvino-cortical feedback projections enhance the net recurrency in the cortical circuit and that this recurrency modulates the evidence accumulation process in the pulvino-cortical circuit.

### Thalamocortical motifs and frequency-dependent interactions

In previous sections we used a computational model to elucidate the role of feedforward and feedback pulvino-cortical pathways in various cognitive behaviors. We now investigate how interactions between these pulvino-cortical pathways are influenced by different network motifs. In Fig. S1 *A*, we sketch three possibilities for the connectivity between two cortical modules and the pulvinar given both types of pulvino-cortical pathways (see also Whalen et al. (2015) for a symmetry-based analysis of three-node networks). First, these pathways could independently coexist so that their function can be deduced from the individual analyses performed so far. Alternatively, these pathways could interact in at least two ways. In one scenario, two distinct pulvinar populations could participate in the feedback thalamocortical and feedforward transthalamic pathways, respectively, and these populations could inhibit each other (see Fig. S1 *B* and Crabtree and Isaac (2002)). This intra-pulvinar competition motif leads to a tradeoff in which one functional circuit is privileged over the other, i.e., a strengthening of a local representation vs propagation of that representation to the next cortical area.

In another scenario, considered broadly in the architecture presented in Fig. 1, the same pulvinar population participates in both feedforward and feedback computations. We now explore this motif in the context of oscillatory processing within and across cortical areas. There is recent evidence from multi-unit activity and local field potentials in the macaque of enhanced coupling between two cortical regions and between cortex and pulvinar at particular frequencies during tasks that engage attention (Saalmann et al., 2012; Zhou et al., 2016). It is not clear what aspect of the connectivity or the dynamics gives rise to the preferential coupling at these frequencies and importantly, how this relates to thalamo-cortical processing. To this end, we reconsidered the multi-regional architecture introduced in Figure 1: two cortical modules (cortical areas 1 and 2) and one thalamic module representing the pulvinar (Figure 7 *A*). As before, the two cortical modules are reciprocally connected via direct cortico-cortical projections and indirectly connected through the pulvinar. To address oscillatory processing in the pulvino-cortical circuit, each of the cortical modules has now laminar structure, in that superficial and deep layers are distinguished on the basis of their connectivity within and across areas. Both layers are composed of excitatory and inhibitory populations that interact to produce noisy rhythmic activity in isolation: superficial layers generate gamma oscillations, while deep layers generate alpha (low beta) oscillations (see Fig.S2 *B* and Mejias et al. (2016)).

In our laminar circuit model the pulvinar module sends feedback projections to the cortical module 1 and relays a transthalamic projection to cortical module 2. After lesioning the pulvinar in our model, we observed an increase in low-frequency oscillations in cortical area 1 (Fig. 7 *B*). We note that feedback connections arising from the thalamus target interneurons in deep layers (Cruikshank et al., 2010; Zhou et al., 2017; Audette et al., 2018). Thus, after a lesion to the pulvinar, pyramidal neurons in the deep layers are disinhibited which subsequently leads to an increase of power in the alpha range due to net excitation in the deep-layer excitatory-inhibitory circuit (Mejias et al., 2016). This result is consistent with the findings by Zhou et al. (2016) who recorded from macaque V4 and observed such increases in alpha-range power after lesioning the pulvinar with muscimol (their figure 7). Indeed, feedback thalamo-cortical projections have been hypothesized to play a modulatory role (Jones, 2007), and in our model they regulate the excitation in the cortical circuit it projects to (Ferguson and Gao, 2017).

We also show that the engagement of the pulvinar via the transthalamic projections enhances the communication at gamma frequencies between the two cortical regions (Fig. 7 *C, D*). Such gamma-mediated coupling is thought to be used for selective communication between cortical areas (Bastos et al., 2015). We computed the spectral coherence between the two cortical areas, which provides a rough estimate of the degree of mutual oscillatory coupling. After lesioning the pulvinar, the spectral coherence between both cortical areas in the gamma range decreases (Fig. 7 *C*), suggesting an important role for the transthalamic connection (see also Fig. S2*A*). These findings are in line with Zhou et al. (2016) who recorded from V4 and IT regions of the visual cortex during a task that required attention and found enhancements in gamma-range coherence (their figures 7 and 8). Interestingly we also found a notable interareal coherence in the alpha range for feedforward communication (see figure 3 from Saalmann et al. (2012) that increases after lesioning the pulvinar. These results extend to Granger causality, which in addition measures directionality: the influence of cortical area 1 to cortical area 2 (2 to 1) is stronger in the gamma (alpha) range and decreases (increases) after a lesion to the pulvinar (Fig. 7 *D*). We propose that the transthalamic projection enhances the transmission of information through a feedforward gamma channel, as increased excitation onto superficial layers in cortical area 2 enhances gamma activity locally (Mejias et al., 2016). This parsimonious interpretation is consistent with our previously described function of pulvinar-mediated modulation of feedforward connectivity: the increased drive from cortical area 1 to cortical area 2 due to the presence or enhancement of pulvinar activity (Figs 3 and 4) is reflected in an increase in gamma oscillations and coherence (Fig. 7). To conclude, the circuit topology presented here instantiates the pulvino-cortical feedforward and feedback pathways concurrently, and we show how the pathways contribute to the frequency-dependent interactions within and across cortical areas.

### The pulvinar can modulate functional hierarchies in the cortex

Spectral Granger causality profiles of cortical interactions like the ones introduced above can be used to define a functional hierarchy, as defined by Bastos et al. (2015). Briefly, two cortical areas C×1 and C×2 are said to show an ascending functional hierarchical relationship if the spectral Granger causality pattern from C×1 to C×2 (C×2 to C×1) is predominantly strong in the gamma (alpha) range. The level of saliency of such pattern can be quantified by the so-called hierarchical distance, which is a function of the Granger causality profiles. Building on the cortical model by Mejias et al. (2016), our laminar model of pulvino-cortical interactions shows the presence of a functional hierarchy between the two cortical areas when the pulvinar is present. After lesioning the pulvinar, we observed a decrease in such hierarchical distance (Fig.7 *D*, inset and Fig.S2*C*). These results suggested that we can obtain a range of hierarchical distances by manipulating the pulvinar gain. As shown in Fig. S3, this is indeed the case: an increase in the pulvinar gain leads to an increase in hierarchical distance between the two cortical modules, consistent with the context-dependent hierarchical jumps observed by Bastos et al. (2015).

Hierarchy can also be defined functionally in terms of the timescale of intrinsic fluctuations during spontaneous activity. Areas high in the cortical hierarchy such as prefrontal areas have larger intrinsic timescales than lower areas such as sensory areas (Murray et al., 2014). We used the 2AFC version of the model in Fig 1 to show that increasing the pulvinar gain gives rise to increasing timescale differences between the two cortical areas (Fig. S3). These results are consistent with a modeling study that proposes that long-range connectivity contributes to the timescales of individual cortical areas (Chaudhuri et al., 2015), but our model goes beyond by suggesting that the pulvinar contributes, in a gain-dependent manner, to these cortical intrinsic timescales. Importantly, the two functional hierarchies we have introduced, i.e., oscillation- and timescale-based, are consistent and are both characterized by an increasing hierarchical distance as a function of pulvinar gain (Fig. S3). We thus propose that the pulvinar contributes to maintaining and modulating the hierarchical relationships between cortical areas.

## Discussion

In this study, we propose a multi-regional circuit model that subserves cognitive computations and is composed of two cortical areas and the pulvinar nucleus of the thalamus. We highlight the functional relevance of two pulvino-cortical pathways: a feedforward pathway that connects two cortical areas transthalamically and a feedback pathway that engages the thalamic reticular nucleus (TRN) and projects back to the cortex. We summarize how the aforementioned pathways contribute to different cognitive computations, including attention, working memory, and confidence during decision-making.

First, lesions to the pulvinar in the model resulted in action-related disruptions in the contralesional field, including increased saccade latency and decreased choice performance in a visuo-spatial task (Desimone et al., 1990; Wilke et al., 2013). These results are consistent with structured cortico-thalamic connections in healthy subjects that allow for distractor-filtering computations during visuo-spatial attention tasks. Second, the circuit model can subserve working memory in the form of spatially-selective persistent activity. Crucially, the pulvinar can switch the pulvino-cortical circuit to subserving a global persistent-activity state as well as establish two different dynamical regimes during distractor processing in working memory. Third, the modulation of the pulvinar can bias the circuit into a predominantly feedforward mode in which bottom-up information is preferentially transmitted as opposed to top-down information. Thus, the pulvinar can induce a cortical network reconfiguration that can be used to resolve decision-making conflict (Rotshtein et al., 2011). Fourth, we suggest that the pulvinar estimates decision-making confidence as a result of plastic cortico-thalamic projections that engage the TRN. Our model provides a unified account of implicit (Kiani and Shadlen, 2009) and explicit (Komura et al., 2013) representations of decision-making confidence in the cortex and pulvinar, respectively. Finally, pulvino-cortical feedforward and feedback pathways can regulate hierarchical frequency-dependent interactions within and across cortical areas. We thus provide a novel and parsimonious interpretation of recent experiments targeting the macaque pulvinar (Saalmann et al., 2012; Zhou et al., 2016).

In light of these modeling results, we suggest that the pulvinar augments the computational capabilities of an otherwise isolated cortical cognitive-type circuit. The cortex “outsources” local and long-range cortical connectivity to the pulvino-cortical feedforward and feedback pathways for an additional layer of control. Indeed, instead of being fixed, pulvino-cortical feedforward and feedback pathways can be dynamically engaged through external modulation of the pulvinar. We propose that such cognitive-circuit outsourcing is an organizational principle for flexible distributed computation in the brain.

### Pulvinar and attentional modulation and deployment

The pulvinar is part of a complex multi-regional circuitry that is involved in attentional processing in humans (Snow et al., 2009; LaBerge and Buchsbaum, 1990; Danziger et al., 2004, 2002; Ward et al., 2002) and non-human primates (Petersen et al., 1985; Desimone et al., 1990; Wilke et al., 2010, 2013; Saalmann et al., 2012; Zhou et al., 2016). Pulvinar cells are modulated by attention (Petersen et al., 1985) and lesions to the pulvinar cause attentional impairments in the contralesional field (Wilke et al. (2010); Snow et al. (2009), see also Karnath et al. (2002) for similar effects from basal ganglia lesions in humans). Attentional processing entails various computations, including spatial shifting and distractor filtering. We suggest that the pulvinar is involved in these computations through reciprocal connections with cortical areas typically recruited in attentional tasks, including the fronto-parietal network (Corbetta and Shulman, 2002; Suzuki and Gottlieb, 2013; Bisley and Goldberg, 2010; Schall, 2015) and superior colliculus (SC) (White et al., 2017). We contend that the filtering ability of the distributed attentional network that includes the pulvinar is intimately related to a connectivity profile where same-selectivity populations excite each other, while opposite-selectivity populations inhibit each other. Particularly in the context of spatial tasks where distractors are located opposite to the target with respect to the meridian, we suggest that interhemispheric competition and inhibition (Szczepanski and Kastner, 2013; Palmer et al., 2012), possibly mediated by the TRN (Viviano and Schneider, 2015), are essential. We predict that compromising these projections through thalamic or TRN lesions are potential sources of the distractor-filtering deficits and hemispatial neglect that are commonly observed in subjects or patients with this type of damage.

Previous models have proposed the existence of a saliency map in the brain that can control the deployment of attention by combining both bottom-up and top-down salience (Itti and Koch, 2001), and the pulvinar may be part of such a map. Interestingly, other thalamic circuits including the LGN and the TRN have been involved in attentional enhancement (Crick, 1984; McAlonan et al., 2008; Wimmer et al., 2015; Halassa and Acsády, 2016). It will be important for future studies to examine and compare contributions from the different thalamic nuclei to computations that generally support selective attention (Buschman and Kastner, 2015; Béhuret et al., 2015).

### Gain modulation through external control of pulvinar excitability

In this study we propose that one key function of the pulvinar is to gate the effective cortico-cortical connectivity via gain modulation (see Cortes and van Vreeswijk (2012) and Olshausen et al. (1993) for proposals for the pulvinar in propagation and routing of information, respectively). Indeed, the pulvinar receives inputs from many structures including the prefrontal cortex (Romanski et al., 1997), the pretectum (Benevento and Standage, 1983), superior colliculus (Baldwin et al., 2013; Berman and Wurtz, 2011; Zhou et al., 2017), and brainstem (Varela, 2014). Assuming that these areas are external to the pulvino-cortical circuit we considered, they could potentially modulate the pulvinar activity as suggested by our model. For example, we found that the gain modulation in the pulvinar could resolve decision-making conflict. We predict that the pulvinar participates in a conflict-resolution system, possibly in conjunction with the cingulate cortex (Botvinick et al., 2004) given that the pulvinar receives connections from it (Romanski et al., 1997). Generally, we suggest that the pulvinar serves as a driver node for open-loop cortical control (Muldoon et al., 2016) (see also Dominguez-Vargas et al. (2017)) that could complement other gating mechanisms relevant for cortical processing (Wang and Yang, 2018).

For simplicity, we have lumped the modulatory effects of the external areas mentioned previously into the control of a single parameter, the pulvinar excitability λ, which in our circuit model represents the slope of the FI curve (Abbott and Chance, 2005). The firing rate vs current (FI) curve and the proposed gain-modulation mechanism in the pulvinar can be further shaped by cortico-thalamic noise (Béhuret et al., 2015) as well as the firing mode of the thalamic relay neurons (Steriade et al., 1990; Saalmann and Kastner, 2011; Sherman and Guillery, 2013). Moreover, other gain-modulation mechanisms may be relevant for selective transmission, including synchronization (Saalmann and Kastner, 2011; Saalmann et al., 2012). We suggest that the gain-modulation of the pulvinar proposed in this model would regulate not only effective cortico-cortical connectivity, but also connectivity between the cortex and other subcortical structures (Zhou et al., 2018).

### Confidence representation in the pulvinar and its relationship to attention

In this modeling study we have examined why and how the pulvinar is involved in confidence in decision-making. These modeling results relate to the study by Komura et al. (2013) who found that the firing rate of macaque pulvinar cells correlated with decision-making confidence during a visuo-spatial categorization task (Komura et al., 2013) (see also Kepecs et al. (2008) for neural correlates of confidence in the rat orbito-frontal cortex). Pertinent to our modeling results, there are implicit signatures of decision-making confidence in the lateral intra parietal cortex (LIP) (Kiani and Shadlen, 2009) (see Teichert et al. (2014) for uncertainty signatures in FEF). We suggest that the confidence representation as observed implicitly in the firing rates of LIP neurons is directly related to the explicit representation of confidence in the pulvinar cells (Komura et al., 2013) (see also Wei and Wang (2015)). Indeed, pulvinar cells estimate confidence by integrating and transforming cortical signals through an absolute-value-type computation (see Fig. 5 *B*, bottom) that involves a plastic cortico-thalamic circuit that engages the TRN. Along these lines we predict that, first, the cortex and pulvinar may be both form part of distributed circuit for decision-making so that lesions or disengagement of the pulvinar causally affect the decision-making process (see speed-accuracy tradeoff in Figure 6 *D, E*) and second, a plastic (and intact) TRN-pulvinar circuit is necessary for the pulvinar to estimate confidence. We note, however, that there are differences in the opt-out task design between the Komura et al. (2013) and Kiani and Shadlen (2009) studies, where the opt-out component is always present in the former, but randomly interleaved in the latter. Future experiments that consider simultaneous recordings of the cortex and pulvinar as well as optogenetic manipulation (e.g., inhibition) of the TRN-pulvinar circuit in the context of a consistent post-decision wagering task could test these predictions.

Here we proposed that the TRN-pulvinar circuit calculates the absolute difference of firing rate activities of two populations from an upstream cortical area. In the framework of predictive coding, such computation might be useful to represent computational precision (Kanai et al., 2015). Furthermore, we suggest that this TRN-pulvinar computation generalizes across tasks and species. For example, Roth et al. (2015) found that the LP, the rodent analogue of the pulvinar, signals the discrepancies, both positive and negative, between self-generated and external visual motion (see in Roth et al. (2015), their figure 7*C*). We suggest that this finding, in this case related to locomotion, is another instance of the canonical computation (Carandini and Heeger, 2012) our plastic TRN-pulvinar circuit can perform. We propose more generally that the pulvinar can represent saliency for visually-related behavior and that this saliency is interpretable as confidence in the case of visuo-spatial decision-making (Komura et al., 2013) or sensory context in the case of behaviors involving locomotion (Roth et al., 2015).

How does the confidence representation in the pulvinar relate to the pulvinar’s involvement in attentional tasks? We found that unilateral lesions to the pulvinar result in an asymmetric gain and connectivity pattern that biases the winner-take-all mechanisms behind visual selection, suggesting that pulvino-cortical input is necessary for normal functioning in this task. On the other hand, in Figure 6 we showed that the pulvinar represents decision-making confidence through a transformation of the incoming cortical activity and importantly, the feedback projection arising from the pulvinar was key in regulating the evidence-accumulation mechanism underlying cortical decision making. We propose that for both attention, i.e., the attentional processes behind distractor filtering, and confidence-related computations, the pulvinar provides contextual modulation to a cortical circuit that processes visual information. Furthermore, we suggest that the computational significance of the signals observed in a given pulvinar region depends on the cortical areas projecting to it. Thus, if areas involved in the decision-making accumulation process (e.g., LIP) project to a particular region of the pulvinar (e.g., medial pulvinar), the signals observed in this region - after an appropriate transformation (Fig. 5) - would be interpreted as decision-making confidence, which can be broadcast to other cortical areas via pulvino-cortical projections. On the other hand, if visual areas earlier along the hierarchy coding for a feature of the visual scene project to a more ventral region of the pulvinar, then the signals observed in this region would be interpreted as visual saliency of that particular feature relative to either other features or to the background (Wilke et al., 2009). Therefore, the signals observed in the pulvinar could reflect behaviorally-relevant transformations of ongoing cortical activity that can be broadcast to other cortical areas.

### The functional and anatomical organization of the pulvinar and other higher-order thalamic nuclei

The pulvinar is endowed with the appropriate circuitry for the computations proposed in this study, namely, open-loop control of the effective connectivity between two cortical areas along the visual pathway and explicit saliency representation within one cortical area. With respect to control of the effective connectivity, the pulvinar is adequate for this computation because its lack of excitatory recurrency results in relatively fast dynamics as compared to the cortex that can aid in the rapid transfer of transthalamic information. Furthermore, the triangular configuration of cortex and thalamus (Theyel et al., 2010) parsimoniously suggests a direct vs indirect means of communication between two areas. Moreover, the pulvinar receives principal projections as well as neuromodulation from a multitude of cortical and subcortical sources including TRN-here referred to as contextual modulation - that can influence the pulvinar activity. Finally, the gross anatomy of thalamo-cortical projections (Shipp, 2015) indicates that visual cortical areas have bidirectional projections to the pulvinar, suggesting that the computations in each cortical area can be modulated via reciprocal loops with a pulvinar buffer (Purushothaman et al., 2012). The pulvinar thus appears to be uniquely positioned to provide contextual modulation to cortical computations associated with cognition as proposed by our model.

The higher-order thalamic nuclei have been less well studied than the first-order sensory nuclei, but there has been recent significant progress on this front. For example, Schmitt et al. (2017) showed that MD thalamic neurons were crucial to maintain task-relevant information during a delay period, but these neurons did not exhibit the rule tuning of its frontal cortical inputs. Analogously, in Fig. 5 we show that differently-tuned cortical populations converge onto the pulvinar so that within the new pulvinar receptive field the dimensionality of the representation decreases (Komura et al., 2013; Schmitt et al., 2017). This organization is different from that of the plots in Figs. 3,4 and of other thalamic nuclei, for which receptive fields tightly reflect their cortical input (Guo et al., 2017; Acsády, 2017). Along these lines, we propose that the pulvinar contains at least two receptive-field types: a receptive field with similar properties to its cortical driving field (for example, Fig. 3), and a receptive field that receives convergent input from differently-tuned cortical populations (see Fig. 5, Schmitt et al. (2017) and figure 8 from Komura et al. (2013)). The different receptive field types in the thalamus might be an organizational principle to define hierarchy in the thalamic system analogous to the structural and functional characterization of hierarchy proposed for the cortex (Felleman and Van Essen, 1991; Markov et al., 2014; Chaudhuri et al., 2015; Shipp, 2015; Mejias et al., 2016; Bastos et al., 2015).

### Model limitations and future directions

Our circuit model can be extended in different ways to address important questions not studied here. For example, we used a simple circuit model to show how pulvino-cortical feedforward and feedback pathways can regulate oscillatory activity within and across cortical areas (Saalmann et al., 2012; Zhou et al., 2016). We note that, although both the Saalmann et al. (2012) and Zhou et al. (2016) studies examined the contribution of the pulvinar to cortico-cortical communication through oscillations, there were important differences that can be addressed in future instantiations of our pulvino-cortical circuit model including task-design, the cortical regions recorded (V4 and TEO, V4 and IT, in Saalmann et al. (2012) and Zhou et al. (2016), respectively) and crucially, the period of the task when the neural data-analysis was performed (stimulus-absent delay period in Saalmann et al. (2012), peristimulus period in Zhou et al. (2016)).

To characterize the local cortical circuit and model 2AFC tasks, we used a parsimonious discrete firing-rate model. A ring model with smoothly-varying tuning would be more appropriate if we wanted to explore the representation of continuous variables such as orientation and/or model multi-item decision making tasks, as well as effects that depend on the distance between distractors and targets for working memory. A spiking circuit with explicit ionic currents such as the low-threshold calcium current would enable modeling the well-documented dual firing modes of thalamic neurons and their participation in thalamo-cortical rhythms (Steriade et al., 1990; Bazhenov et al., 2002). Furthermore, an investigation into the dynamics of ionotropic and metabotropic receptors and their respective timescales could refine the hypotheses concerning the function of different thalamo-cortical pathways as introduced here (Sherman and Guillery, 2013; Sherman, 2016). Finally, an extended version of the thalamo-cortical circuit would include other areas such as the basal ganglia to study, for example, gating of visuo-spatial working memory (Cohen and Frank, 2009) and inhibitory control (Wei and Wang, 2016).

In this computational study we proposed a circuit model to study pulvinar computations in the context of behaviorally-relevant representations in the cortex. The engagement of feedforward and feedback pulvino-cortical pathways constitutes a paradigmatic example of computations in the brain underlying flexible behavior and control. Our interpretation of the function of feedforward and feedback thalamocortical loops offers a novel perspective on cortico-subcortical processing in general and, moreover, will provide solid ground for the development of large-scale models of the brain (Chaudhuri et al., 2015; Mejias et al., 2016; Joglekar et al., 2018) that incorporate the thalamus in dynamical interplay with the cortex.

## Methods

### Model architecture

We constructed a distributed circuit model that is comprised of two reciprocally interacting cortical modules as well as a thalamic (pulvinar) module (Fig. 1). Each module contains two selective, excitatory populations, labeled *A* and *B*. In the mean-field description we consider here, the activity of each population is described by a single dynamical variable (see *Cortical and thalamic circuit dynamics* for details). Within the cortical modules, the two populations have recurrent excitatory connections and interact through a local inhibitory population (not explicit in the Fig. 1 schematic) that allows for cross-inhibition between the two excitatory populations. Each recurrently-connected excitatory population receives inhibition from another population representing a common pool of interneurons. Inhibition is linearized so that projections between the two excitatory populations *A* and *B* are effectively represented by negative weights (Wong and Wang, 2006). The two cortical modules interact through long-range projections that are structured according to the stimulus selectivity of populations within each module, i.e., populations with the same selectivity are connected through excitatory projections whereas populations with different selectivity are connected via net inhibitory projections. This configuration allows the circuit to subserve winner-take-all competition, slow integration for decision making, as well as to maintain stimulus-selective persistent activity (Wong and Wang, 2006; Wong et al., 2007; Murray et al., 2017).

The pulvinar module also contains two excitatory populations. However, the excitatory populations do not interact through locally-recurrent excitatory projections (Jones, 2007). The thalamic populations can, however, interact via local interneurons (as in the medial pulvinar of the primate (Imura and Rockland, 2006)) or through interactions with the inhibitory cells of the thalamic reticular nucleus (TRN). The cortical modules are connected with the thalamic module through cortico-thalamic feedforward and feedback pathways (Sherman and Guillery, 2013). The cortico-thalamic feedforward - or transthalamic - pathway refers to projections from one cortical area to the thalamus, and these projections are relayed to a second cortical area (Sherman, 2016). In our model the transthalamic projections are topographic as in the cortico-cortical connections: same-selectivity populations are connected through excitatory projections while opposite-selectivity are connected through inhibitory projections. The pulvino-cortical feedback pathway refers to connections between one cortical area and the pulvinar that are reciprocated to the same cortical area. These connections include a cortical monosynaptic excitatory as well as a disynaptic inhibitory projection through the TRN. In our model we consider concurrent pathways, i.e., the pulvinar module participates in both pathways as in Figure 1, but in Fig. S1 we consider other interaction motifs. In the section *Connectivity* we formalize these assumptions with specific values for each of the connections.

### Cortical and thalamic circuit dynamics

We first consider the dynamics of neural populations in the cortical modules. Each cortical population *i* = *A, B* is described by one dynamical variable, its average firing rate. The firing-rate dynamics of the population *i* in the cortical modules are dominated by the slow dynamics of the average NMDA synaptic gating variable *s*_*i*_. Indeed, the dynamics of the NMDA synaptic gating variable is slow compared to the other time scales in the system so that the other dynamic variables, i.e., GABA and AMPA gating variables, are described by their steady state values (Wang, 2002; Wong and Wang, 2006; Wong et al., 2007; Murray et al., 2017). The dynamical equation for the NMDA gating variable *s*_*i*_ for the cortical module *n* = 1, 2 is:

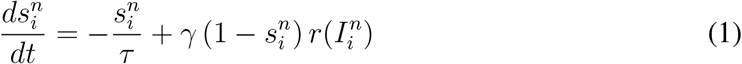

where τ = 60 ms is the NMDA time constant, γ = 0.641 controls the rate of saturation of *s*, and *r*(*I*_*i*_) is the firing rate of the population *i* as a function of the input current *I*_*i*_. The firing rate as a function of input current is given by the frequency-current (F-I) curve relation (Abbott and Chance, 2005,

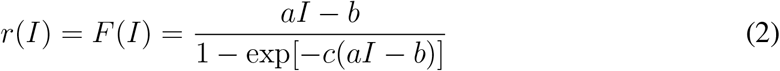

with 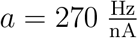, *b* = 108 Hz, and *c* = 0.154 s.

For the two neural populations in the pulvinar module *p*, we also consider a one-variable dynamical equation for each population. In the circuit model, the non-recurrent dynamics in the thalamo-cortical relay cells are mediated primarily by non-NMDA currents (Golshani et al., 1998; Bazhenov et al., 2002) so that the dynamical equation for the thalamic gating variable 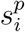, *i* = *A, B* is:

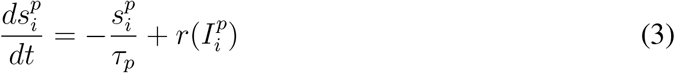

where *τ*_*p*_ = 2 ms is time constant of fast AMPA thalamo-cortical synapses and 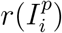 is the firing rate of the pulvinar cell population *i* as a function of the input current *I*_*i*_. As in the cortical modules, the thalamic firing rate as a function of input current is given by the frequency-current (F-I) curve relation (Abbott and Chance, 2005):

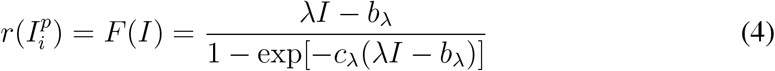

where λ is the pulvinar F-I slope, here referred to as the pulvinar excitability (the value of λ lies between 120 and 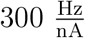 and is reported in the figure captions), *b*_λ_ = 112 Hz, and *c*_λ_ = 0.2 s. The values chosen result in realistic firing rates for pulvinar neurons (Dominguez-Vargas et al., 2017; Komura et al., 2013).

The input current to population *i* = *A, B* in both cortical modules is given by:

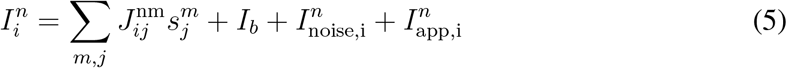

where the first term of the right-hand side of Eq. 5 corresponds to synaptic inputs from cortex and thalamus: 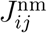 is the connection weight from population *j* in Module *m* = 1, 2,*p* to population *i* in cortical Module *n* =1, 2, *I*_*b*_ is the background current, 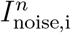 is the noise current to population *i* in Module *n*, and 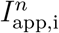 is the applied current to population *i* in Module *n* from external sources. Below we describe the noise and applied currents in detail. Similarly, the input current to population *i* = *A, B* in the pulvinar is given by:

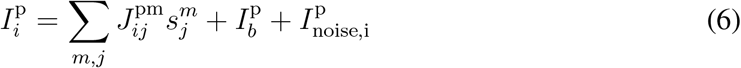

where 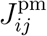 is the connection weight from population *j* in the cortical Module *m* to popudlation *i* in Module *p*, 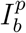 is the background current,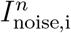 is the noise current to population *i* in Module *p*, and 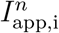 is the applied current to population *i* in Module *n* from external sources, typically bottom-up (sensory) or top-down (internal).

For the cortical and thalamic modules, we mimic external non-selective currents through a noise current to each population. The noise current follows Ornstein-Uhlenbeck dynamics with the time constant of AMPA synapses:

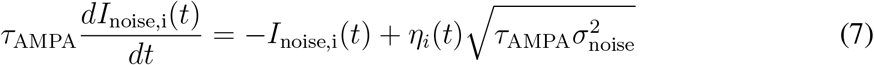

where *τ*_AMPA_ = 2 ms, *η* is Gaussian white noise with zero mean and unit variance, and σ_noise_ sets the strength of noise. Parameter values are reported in Table 1.

We consider the external current *I*_app_ to the cortex for the following scenarios: i) Visual selection (Wilke et al., 2010, 2013; Dominguez-Vargas et al., 2017; Desimone et al., 1990), ii) working memory and distractors (Suzuki and Gottlieb, 2013), iii) decision-making and confidence (Kiani and Shadlen, 2009; Komura et al., 2013). We will specify these external currents after the Connectivity section below.

### Connectivity

The connectivity in our model is specified by the sign and magnitude of the connection weights between the selective excitatory populations for each of the three modules: two cortical, one thalamic (pulvinar). We first specify the connectivity for the two-module cortical model (for additional details see Murray et al. (2017)). The connections can be local (within a module) and long-range (across modules). To this end, it is useful to express the connection weights with the terms:

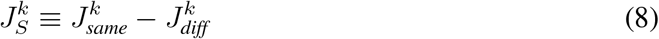

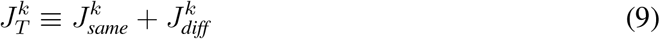

where *J*_*same*_ denotes the positive connection weight between same-selectivity populations, e.g. from population A in Module 1 to population A in (cortical) Module 1 or 2. *J*_*diff*_ denotes the negative connection weight between different-selectivity populations, e.g. from population A in Module 1 to population B in Module 1 or 2, and *k* = 11,12, 21, 22 defines whether the connection is local or long range. We define *J*_*S*_ as the *structure* of the network, since it reflects the magnitude of same-selectivity excitation and different-selectivity cross-inhibition and thus the total recurrent strength. Analogously, we define *J*_*T*_ as the *tone* of the network, which reflects the net input onto a particular population. For both long-range projections between modules, we constrain them to have pathway-specific excitation/inhibition (E/I) balance:

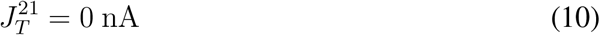

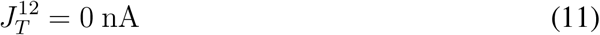

We can easily translate the structure *J*_*S*_ and tone *J*_*T*_ into individual synaptic weights. For example, 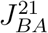 denotes the feedforward projection between the population *A* in the first module onto the population *B* in the second module and is given by:

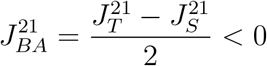

We now describe the connectivity between the cortical modules and the pulvinar. Cortico-thalamic projections *J*^*pk*^ that target the pulvinar are represented by matrices of the form

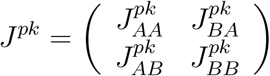

where *k* = 1, 2 are indices of the cortical modules and *A, B* denote the stimulus selectivity. Thus, 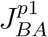, for example, represents the inhibitory weight between population *A* in module 1 and population *B* in the pulvinar. Furthermore, connections are symmetric in that 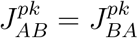 and 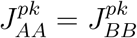.

Pulvino-cortical projections *J*^*kp*^ that target cortical Modules 1 and 2 are analogously represented by matrices of the form

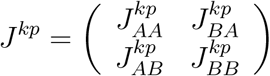

where as before, *k* =1, 2 are indices of the cortical modules and *A, B* denote the stimulus selectivity. For both cortico-thalamic and thalamo-cortical excitatory projections we define a generic excitatory projection *J*_exc_ = *w* · *b*_*p*_, where *b*_*p*_ is a baseline value and *w* ∈ {*w*_1*p*_,*w*_*p*1_,*w*_*p*2_,*w*_2*p*_} determine the connection weights. For example, 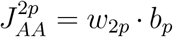 denotes the excitatory connection strength between the A population in the pulvinar and the *A* population in the cortical module 2. For both cortico-thalamic and thalamo-cortical inhibitory projections, we define a generic inhibitory projection as *J*_inh_ = *c*_inh_ · *J*_exc_ where *c*_inh_ dictates the degree of excitatory-inhibitory balance for that pathway. *c*_inh_ = −1 implies full balance in that *J*_exc_ + *J*_inh_ = 0. For example, and in the case of full balance, 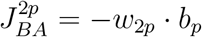 denotes the inhibitory connection strength between the *A* population in the pulvinar and the *B* population in the cortical module 2. Thus, the connectivity between the cortical modules and pulvinar in our circuit model is completely specified by assigning values to the cortico-thalamic (and thalamo-cortical) projection parameters *w*,*b*_*p*_, and *c*_inh_. Since the anatomical data to fully specify the values for these projection parameters in this framework is not available (but see Oh et al. (2014)), we used the following general constraints: the total cortico-thalamic projection weight is greater than the total thalamocortical weight (Jones, 2007), the feedforward relay weights in the hierarchy-preserving direction (cortical area 1 - pulvinar - cortical area 2) is larger than in the reverse direction (cortical area 2 - pulvinar - cortical area 1, see Sherman (2016)). The values for the projection parameters are in Table 1.

Given the input currents to thalamic and cortical cells specified by Eqs. 5 and 6 and the connectivity specified above, we can now write the general pulvino-cortical model as

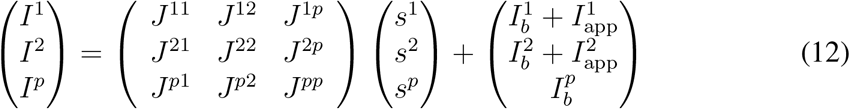

where *I*_*n*_ (*n* =1, 2,*p*) is the total current in each Module *n*, *J*^*mn*^ are synaptic weight matrices connecting modules *n* to *m*, *s*^*n*^ are the corresponding synaptic gating vectors, and 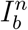 and 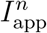 are base and applied input currents, respectively.

### Visual selection and pulvinar lesions

In Figure 2 we simulated a decision-making task (Wilke et al., 2013) analogous to target selection during visual search (Schall, 2015). Each module contains two populations that are selective to a target and a distractor, respectively. A distractor was defined as another stimulus simultaneously flashed at an opposite location to the stimulus (Desimone et al., 1990). External stimuli enter as currents into the cortical modules. The external currents are segregated into “bottom-up” corresponding to sensory-type inputs and “top-down” inputs, corresponding to reward expectation, task representations and/or working memory. These applied currents reflect the external stimulus as:

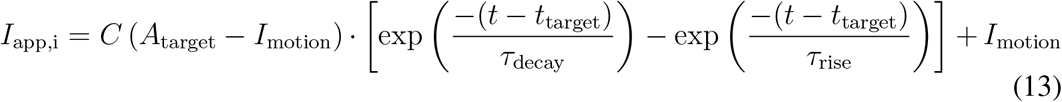

where the first term on the right-hand side represents the transient to the visual stimulus, 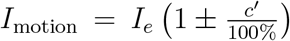 represents the sensory evidence, *I*_*e*_ scales the overall strength of the input and *c′*, referred to as the differential input, sets the bias of the input for one population over the other (equivalent to the coherence in Wong and Wang (2006)) and represents the target-distractor similarity in a visual search task, *A*_target_ and *t*_target_ determine the amplitude and the onset of the target, respectively, the time constants *τ*_decay_ and *τ*_rise_ determine approximately the decay and rise of the target-induced transient response, and *C* is a normalization factor (see values in Table 1). In our model, the target is associated with a larger value of the differential input *c*′. A zero-c’ stimulus applies equal input *I*_*e*_ to each population in Module 1. In all of the simulations and when *c*′ > 0, the target-selective population receives the greater biased input. Due to noise, however, this does not guarantee that the target population will win, especially for low c’ values. Finally, we modeled a lesion by setting the firing-rate of one of the pulvinar populations to zero.

We modeled reward expectation in the task (Wilke et al., 2013) as a current *I*_reward_ applied to Module 2, modeled as in Eq. 13 but without the visual transient. A saccade to a particular direction was defined as the action obtained after a population selective to a stimulus at that location reaches the firing-rate threshold of 30 Hz. Saccade latency was measured as the time at which the firing rate of a population in Module 2 crossed the 30 Hz threshold. For the Choice experiment in Fig. 2, we varied the reward-expectation current onto Module 2 in the scenario lesion+reward so that the new *I*_reward_ = 0.012 nA. Proportion of saccades in the Choice task was measured as the fraction of saccades made to either side given the differential input *c*′ = 0. For the distractor-filtering simulations in Figure 2 *C*, we simulated lesions in the “target” and “distractor” scenarios, where the lesion was on the pulvinar population selective to the target and distractor, respectively.

### Gain modulation and effective cortico-cortical connectivity

The system of equations that describe the dynamics in the pulvino-cortical circuit is given by Eq.12. We will find an approximately equivalent reduced system. To this end, we make the following two approximations. First, we assume that the synaptic dynamics in the pulvinar are much faster than the dynamics in the cortical loop model (τ_*p*_ ≪ τ_*s*_) and second, we approximate the FI curve in Eq. 4 as

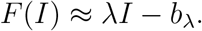

We can then write a reduced description of Eq.12 as

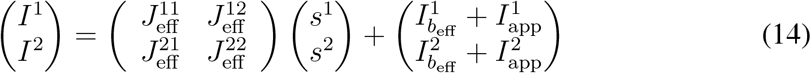

where the effective connectivity matrices are

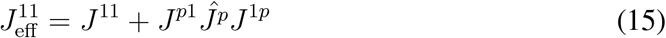

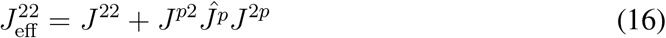

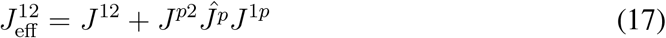

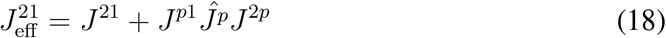

where

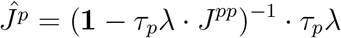

and *J*^*pp*^ describes the interactions within the pulvinar and TRN. Moreover, the new effective base currents are

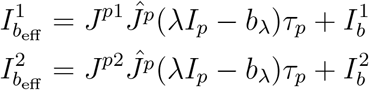

We can write the effective long-range structure 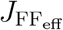 (recurrent excitation and cross-inhibition, Murray et al. (2017)) in the feedforward direction as

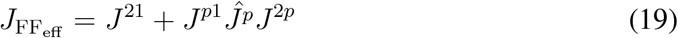

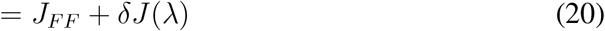

where *J*^21^ ≡ *J*_*FF*_ is the original, i.e., anatomical, feedforward structure from Modules 1 to 2, and δ*J*(λ) is the pulvinar-excitability-dependent transthalamic weight (see Figure 4*A*). As mentioned before, we assume that the feedforward cortico-thalamic and thalamocortical weights in the hierarchy-preserving direction (cortical area 1 - pulvinar - cortical area 2) are larger than in the reverse direction (cortical area 2 - pulvinar - cortical area 1, see Sherman (2016)). We can then write the ratio of feedforward-to-feedback connectivity

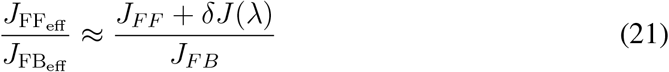

### Simulation of a working memory task with and without distractors

For Fig. 3 we simulated two versions of a working memory task. In the first task, the subject must remember the location of the stimulus across a delay period. A flash of 100 ms appears on one of two positions of a screen indicating the target position. In the second version of the task, a distractor is presented during the delay period after 800 ms. The target to be held in WM is the first stimulus presented. We set the target as a current *I*_app,A_ = *I*_target_ of 100-ms duration that is applied to population *A* in the Module1. Distractors are defined as inputs *I*_app,B_ = *I*_distractor_ of equal duration applied to population *B* arriving after the target and at an opposite location of the visual field. An error (or the “remember-last” regime) is recorded when the population selective to the distractor is at the high memory state at *t* > 3000 ms. We considered three values of the pulvinar excitability λ: small (corresponding to ‘off’, λ = 120 Hz/nA), moderate (λ = 220 Hz/nA) and large (λ = 290 Hz/nA).

### Simulation of a decision-making task with conflicting choices

In Fig. 4, we simulated a decision-making task where bottom-up signals or processes were in conflict with top-down signals or processes. More precisely, we simulated a two-alternative forced choice task in two scenarios: a congruent scenario, in which both bottom-up and top-down currents favored the blue target and a conflict scenario, in which the bottom-up signal favored the blue target in cortical area 1, while the top-down signal favored the red target in cortical area 2. We considered two values of the pulvinar excitability λ: small (λ = 220 Hz/nA) and large (λ = 280 Hz/nA).

We calculated the probability of cortical switching during conflict, i.e., the probability that cortical area 1 enforces its preferred selectivity onto cortical area 2 (“Prob Cx1 switches Cx2”, y-axis in Figure 4*C*), as a function of pulvinar gain λ and conflict level. We calculated the fraction of instances (250 trials in total) where the blue population (favored in cortical area 1) won the competition while parametrically varying λ from 220300 Hz/nA. The high (low) conflict level is given by *c*′ = 20(10).

### Plastic cortico-thalamic projections and confidence representation

We propose a circuit model composed of a cortical module and the pulvinar to elucidate the mechanisms behind confidence-related computations in cortex and thalamus. We first characterize the cortico-thalamic projections in detail and in particular, include short-term plasticity dynamics (Crandall et al., 2015). The schematic of the circuit is shown in Figure 5. We consider two cortical populations that project to the pulvinar. A cortical population projects directly to the pulvinar forming an excitatory synapse and also indirectly through the TRN, forming an inhibitory synapse. Importantly, both the excitatory and inhibitory connections arise from the same cortico-thalamic projection. The excitatory cortico-thalamic synapse exhibits short-term facilitation so that the dynamics of the respective gating variable *s*_exc_ are:

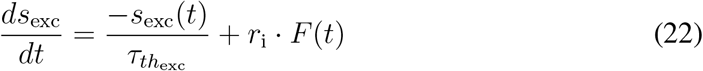

where 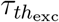 is the time constant of cortico-thalamic excitation, *r*_i_ (i = *A, B*) is the cortical firing rate, and *F* is the facilitation dynamic variable with equation:

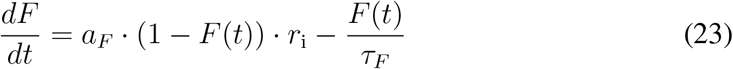

where *a*_*F*_ determines the amount of facilitation and *τ*_*F*_ is the facilitation time constant. The TRN is connected to the pulvinar via an inhibitory synapse that exhibits short-term depression:

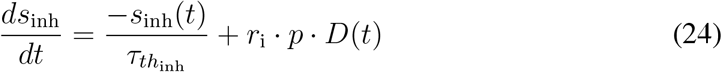

where 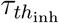 is the time constant of reticulo-thalamic inhibition, *r*_i_ is the cortical firing rate, *p* is the synaptic release probability, and *D* is the depression dynamic variable with equation:

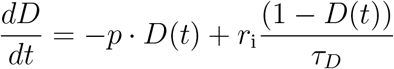

where *τ*_*D*_ is the timescale of depression. Using Eqs. 22 and 24, we can write the total current *I*_i→p_ from a cortical population with firing rate *r*_*i*_ (*i* = *A, B*) to the pulvinar a sum of excitatory and inhibitory components:

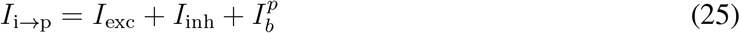

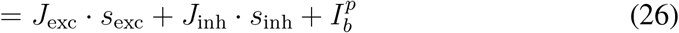

where 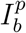 is an external noisy base current to the pulvinar. The steady-state value of the pulvinar firing rate as a function of cortical firing rate is shown in Fig. 5 *B*, top. The current calculated in Eq.26 is for a given cortical population firing rate. In the context of decision making, two cortical populations integrate sensory evidence and compete in a winner-take-all fashion. The total current is thus a contribution from both cortical populations, i.e.,

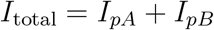

Finally, the firing rate of the pulvinar is, as before, a non-linear function of the current:

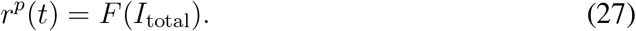

After obtaining the firing rate as a function of the current, we can calculate the steady-state firing rate of the pulvinar as a function of the difference in firing-rate activities at the level of the cortex, shown in Fig. 5 *B*, bottom. The plot resembles a scaled absolute-value function in that the pulvinar activity in the y-axis, which we call “estimated difference” is a symmetric and positive function of the difference in cortical activities, which we call “real difference”. The pulvinar thus performs an approximate absolute-value calculation of the difference between cortical activities (see an intuitive description of this calculation in the Results section).

We modeled perceptual decision-making with an opt-out component in Fig. 6*B*: the subject has the option to forgo the decision and opt for a smaller reward (Kiani and Shadlen, 2009; Komura et al., 2013). The subject must integrate evidence to decide between two orthogonal motion directions, *A* and *B*, corresponding to up and down, for example (Komura et al., 2013). The strength of sensory evidence is modeled as an external current to the two populations as

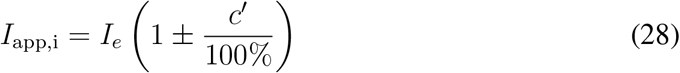

where *I*_*e*_ = 0.007 nA scales the overall strength of the input and *c*′, referred to as the differential input, sets the bias of the input for one population over the other (equivalent to the coherence in Wong and Wang (2006)). For direct comparison with the results from (Komura et al., 2013), we mapped *c*′ to the related measure ‘up-down ratio’ as

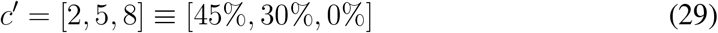

where [45%, 30%, 0%] represents the fraction of dots in the ‘up’ direction (with [55 %, 70%, 100%] representing the fraction of dots in the ‘down’ direction). Easy trials thus correspond to *c*′ = 8 ≡ 0% or 8 ≡ 100%, medium to *c*′ = 5 ≡ 30% or 5 ≡ 70%, and hard to c′ = 2 ≡ 45% or 2 ≡ 55%.

A decision in the model was recorded at a predefined decision time *d*_*T*_ (Kiani and Shadlen, 2009). A correct trial is recorded if the firing-rate activity *r*_*A*_ > *r*_*B*_ and |*r*_*A*_ − *r*_*B*_| > ∊. (Compare to Wei and Wang (2015), who used another population selective for the opt-out target). Analogously, an error trial is recorded if the firing-rate activity *r*_*B*_ > *r*_*A*_ and |*r*_*A*_ − *r*_*B*_| > ∊. Finally an escape trial is registered when at the decision time *d*_*T*_, |*r*_*A*_ − *r*_*B*_| < ∊. For the normalized activities plot in Fig. 5*B* (bottom-right), we calculated the average firing rate in a 250 ms window before the decision time. In the reaction-time (RT) version of the task, we define and calculate the RT as the time at which the firing rate of a population crosses a predefined threshold of 30 Hz. For the RT task, we used a standard set of coherence values to characterize the sensory input (Roitman and Shadlen, 2002): c’ = [0, 3.2, 6.4, 12.8, 25.6].

### Pulvino-cortical circuit with laminar structure

Here we describe the cortico-pulvinar model with laminar cortical structure used in Fig.7. This model extends our previous computational multi-scale framework (Mejias et al., 2016) by introducing a pulvinar module and connecting it to and from cortical laminar populations according to anatomical evidence (e.g. Sherman and Guillery, 2013; Jones, 2007). For simplicity, we have considered only one pulvinar module and up to two cortical populations, but generalizations can be made to accommodate larger thalamocortical networks.

#### Laminar cortical circuit

The circuit of a cortical area consists in two interconnected laminar modules, one corresponding to supragranular (layer 2/3) neurons and another to infragranular (layer 5/6) neurons. Each laminar module contains a recurrently connected excitatory and inhibitory population, with dynamics described by Wilson-Cowan dynamics. The firing rate dynamics of all four populations of a cortical area are given by

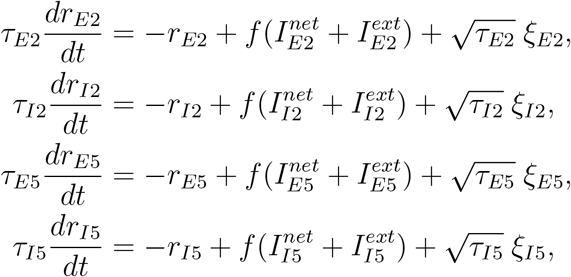

where *r*_*E*2,*I*2,*E*5,*I*5_ are the mean firing rates of the excitatory and inhibitory populations in supra- and infragranular layers, respectively. The corresponding time constants, denoted by τ, are 6,15, 30 and 75 ms. ξ ≡ ξ(*t*) are Gaussian white noise terms of strength *σ* (of values 0.3, 0.3, 0.45, 0.45 respectively), and *f*(*x*) = *x*/(1 − *e*^−*x*^) is the transduction function, or f-I curve, of the neurons. The network input, *I*^*net*^, is the input arriving to each population from other populations in the network -from the same layer, a different layer, or different areas. The terms *I*^*ext*^ are the input from external sources such as sensory stimuli or areas not explicitly included in the model.

Taking into account only local contributions (i.e. assuming an isolated cortical area) the network input is given by

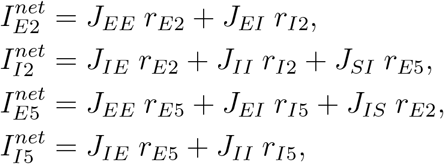

where *J*_*αβ*_ is the mean synaptic strength from population *β* to population *α*. Indices *E*, *I* refer to the excitatory and inhibitory populations of the same layer, and the inter-laminar projections are denoted as *J*_*SI*_ and *J*_*IS*_. Parameter values are *J*_*EE*_ = 1.5, *J*_*IE*_ = 3.5, *J*_*EI*_ = −3.25, *J*_*II*_ = −2.5, *σ*_*E,I*_ = 0.3, *J*_*IS*_ = 1 and *J*_*SI*_ = 0.75. With these parameter values, the circuit displays irregular, noise-driven oscillations in the gamma (supragranular) or alpha (infragranular) rhythms.

*Pulvinar module* To extend our local cortical circuit and include interactions with the pulvinar, we consider a population of excitatory pulvinar neurons of firing rate *r*_*p*_ governed by the following dynamics:

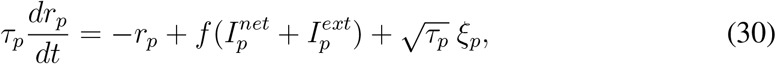

with time constant *τ*_*p*_ = 6 ms, and Gaussian noise *ξ*_*p*_ of strength *σ* = 0.75. The pulvinar population receives input from pyramidal layer 5/6 cells, 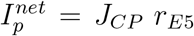, with *J*_*CP*_ = 0.5. Pulvinar also projects back to all cortical populations *E*2,*I*2,*E*5,*I*5, with projection strengths 0.15, 0.1, 0.05 and 0.65, respectively.

### Hierarchical cortico-pulvinar model

To consider how cortico-cortical interactions are modulated by pulvinar activity, we introduce a second cortical area, assumed to be higher in the cortical hierarchy than the first one. Following (Mejias et al., 2016), we consider a feedforward cortico-cortical projection from *E*2 in the first area to *E*2 in the second area, with projection strength *J*_*FF*_ = 1. In addition, we modeled a pulvinar contribution to the feedforward interaction via a pulvino-cortical projection to *E*2 in cortical area 2, with projection strength 0.5. Finally, cortico-cortical feedback projections stem from *E*5 in cortical area 2 and reach *E*2,*I*2,*E*5,*I*5 in cortical area 1 with strengths 0.1, 0.5, 0.9 and 0.5, respectively. External input to all excitatory cortical populations (both laminar modules in cortical areas 1 and 2) and pulvinar is *I*_*ext*_ = 8.

The simulated LFP *R* used to estimate the spectral coherence and Granger causality interactions in the model, is estimated as *R* =(1 − *η*) *r*_*E*2_ + *η r*_*E*5_ with *η* = 0.85, so that both layers contribute to the field signal (although in different ways, given that layer 5/6 pyramidal cells are generally larger). To compute the spectral pairwise conditional Granger causality (GC) between the two cortical areas, we use the Multi-Variable Granger Causality Toolbox (Barnett and Seth, 2014) with an optimal AIC model order of up to 120 ms. We compute the functional hierarchical distance between cortical areas (Fig. 7, S2, and S3) by following the procedure in Bastos et al. (2015) and Mejias et al. (2016). Briefly, we define the directed asymmetry index (DAI) between two cortical areas as the normalized difference between (GC) measurements in both directions, or

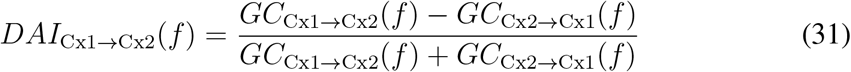

We obtain the multi-frequency DAI index (or mDAI) between two areas by averaging their DAI at the gamma and alpha ranges (and flipping the sign of the alpha term), or

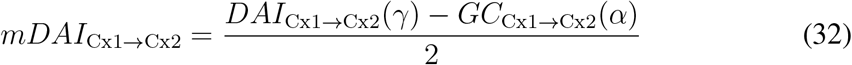

We consider the gamma range as [30, 70] Hz, and the alpha/low beta range as [6, 18] Hz. Since in the present study we only consider two cortical areas, the value of mDAI for this pair gives the oscillation-based hierarchical distance between them. To calculate the oscillation-based hierarchical distance as a function of pulvinar gain (Fig. S3), we varied the (normalized) pulvinar gain *k* as

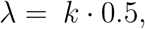

where *k* = 0,1,2,3,4,5.

We also characterized functional hierarchies in terms of intrinsic timescales (Murray et al., 2014). We performed an autocorrelation on the firing rate calculated during spontaneous activity (only noise as input) to reveal the intrinsic or fluctuation time scales of spontaneous activity. The firing rate was first filtered with a Gaussian function with window σ_fiiter_ = 20 ms. To compute the autocorrelation of the firing rate, we substracted the mean from the firing rate and then normalized. We then used Equation 33 to fit the normalized firing-rate autocorrelation and extract the intrinsic time scale τ_int_ as

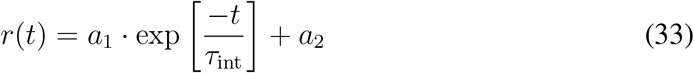

where τ_int_ is the intrinsic timescale during spontaneous activity and *a*_1_ and *a*_2_ are parameters of the fit. The intrinsic timescale difference in Fig. S3 was calculated as the intrinsic timescale in cortical area 2 minus the intrinsic timescale in cortical area 1. We calculated timescale differences as a function of pulvinar gain to produce the plot in Fig. S3, by varying the (normalized) pulvinar gain *k* as

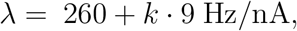

where *k* = 0,1,2,3,4,5.

---

**Figure S1:**
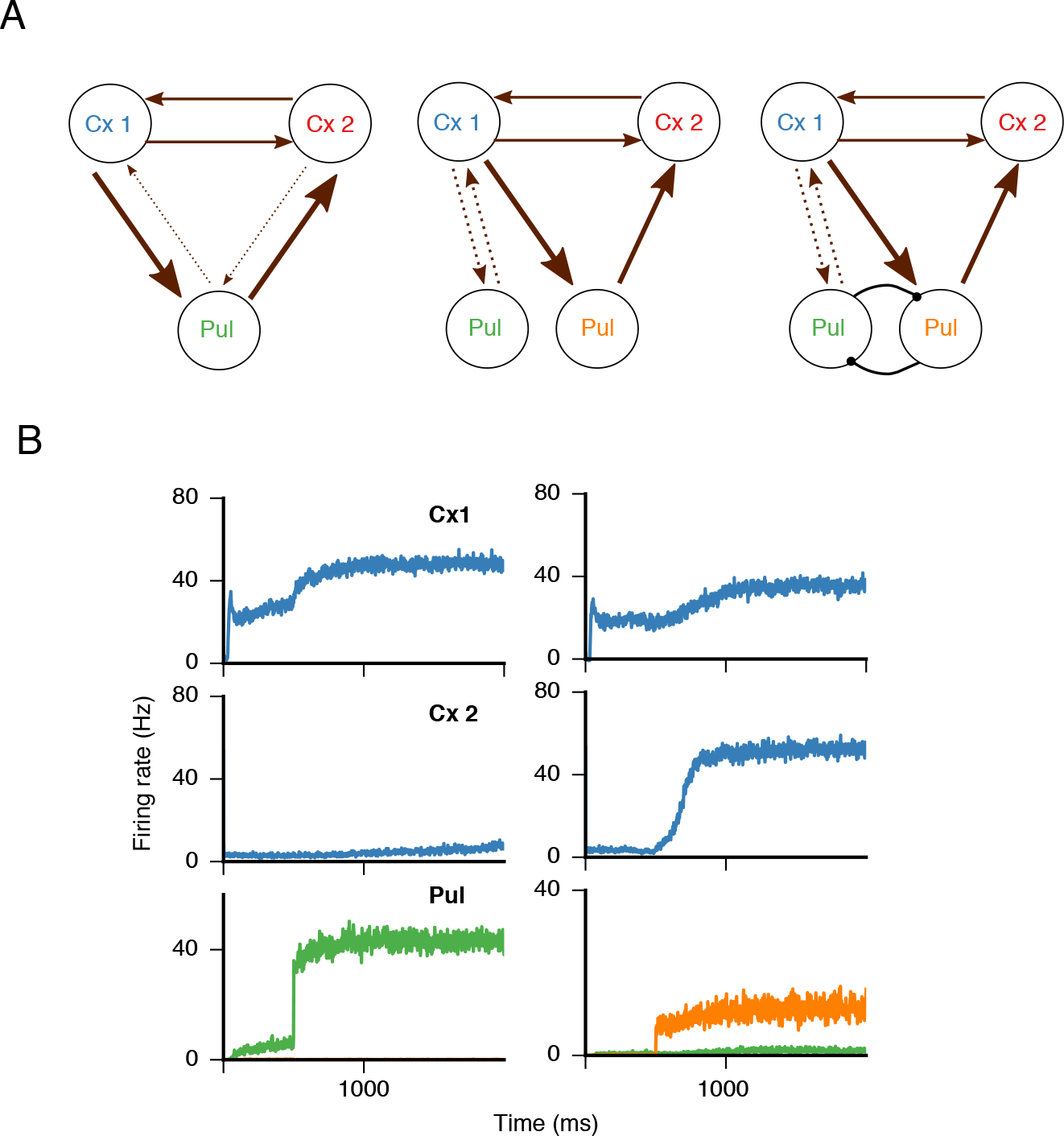
(Related to Fig. 1) Pulvino-cortical pathways interact within different thalamocortical motifs. ***A,*** Three possible architectures connecting pulvinar to cortex: concurrent (left),independent (center), competing (right). ***B,*** For the “competing” architecture in ***A,***, right, there is a tradeoff depending on which of the two pulvinar populations, green or orange, is activated: either a strengthening of a local cortical representation (left, green population active) or propagation of that representation to the next cortical area (right, orange population active). If the strength of recurrent connections is a proxy for noise correlation structure (Helias et al., 2014), this intra-pulvinar competition scenario is consistent with a increase (decrease) of noise correlations across (within) cortical areas as a result of attention (Ruff and Cohen, 2016), here modeled as a bias to a pulvinar population. Furthermore, we suggest that the cortical area whose representation was strengthened will have precedence over the control of subcortical motor centers via projections from layer V axons. This proposal is distinct from - although not necessarily incompatible with - another hypothesis of motor control whereby corticothalamic projections arising from layer V are efference copies relayed to higher cortical areas to monitor impending actions (Sherman and Guillery, 2013; Sherman, 2016)

**Figure S2:**
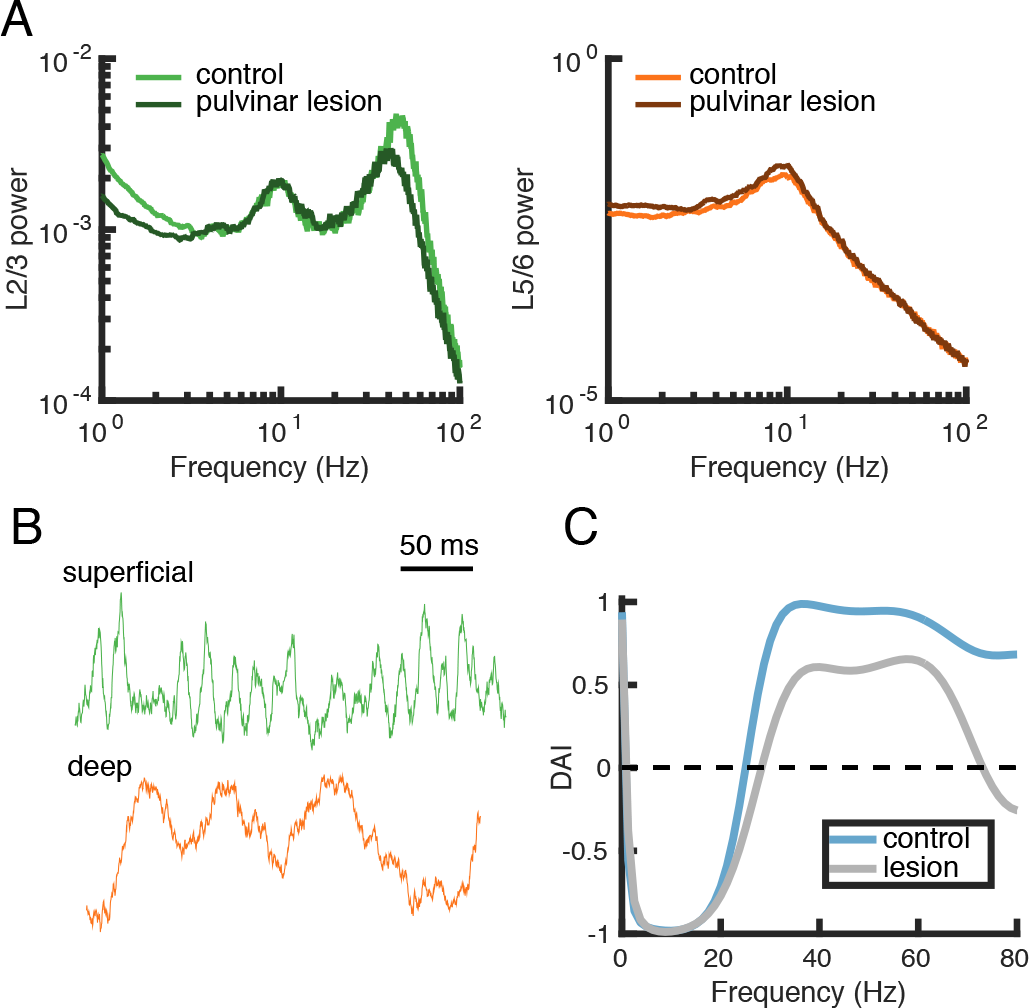
(Related to Fig. 7) Temporal and spectral profiles of the pulvino-cortical circuit before and after pulvinar lesions. ***A,*** Power spectrum for superficial (left) and deep (right) layers for cortical area 1 in control and pulvinar-lesion conditions. ***B,*** Example oscillatory firing-rate traces for superficial and deep layers in cortical area 1. ***C,*** The directed asymmetry index (DAI) for the functional connections between cortical area 1 and cortical area 2 is obtained by normalizing Granger-causality profiles in Fig. 7*D*.

**Figure S3:**
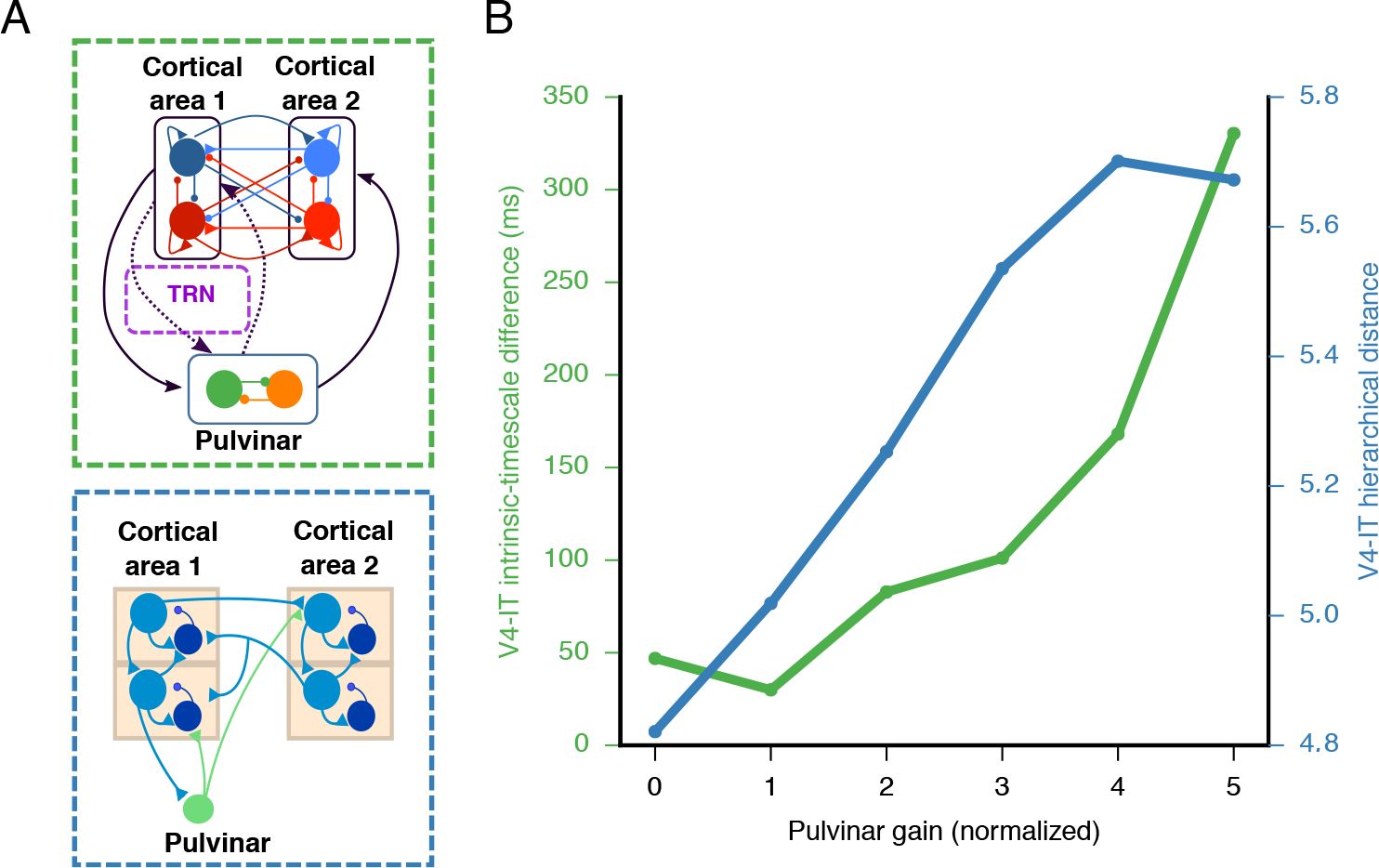
(Related to Fig.3,4, and 7) Pulvinar gain modulates hierarchical distance between two cortical areas. ***A,*** Two instantiations of the thalamocortical model: non-linear model for 2AFC tasks (top, see also Fig. 1) and laminar model for oscillatory coupling between areas (bottom) ***B,*** Both intrinsic timescale difference (green) as well as oscillation-based hierarchical distance (blue) increase as a function of pulvinar gain.

